# Engineering autologous ossicles for the personalized modeling of acute myeloid leukemia

**DOI:** 10.64898/2026.04.13.715606

**Authors:** Camille Sauter, Dimitra Zacharaki, Alejandro Garcia Garcia, David Hidalgo Gil, Yuan Li, Ellen Juel Pørtner, Ani Grigoryan, Medea Gabriel, Aurélie Baudet, Sara González Antón, Adam Cohen Simonsen, Jörg Cammenga, Vladimir Lazarevic, Paul E Bourgine

## Abstract

Acute Myeloid Leukemia (AML) is the most lethal hematological malignancy in adults and is characterized by significant genetic and cytogenetic heterogeneity. This diversity drives substantial patient-to-patient variability in disease progression and therapeutic response. Increasing evidence indicates that bone marrow mesenchymal stem/stromal cells (MSCs) play a critical role in AML emergence, evolution and treatment resistance. However, progress in the field is challenged by the lack of experimental models supporting sustained AML survival while faithfully recapitulating the human bone marrow niche. Humanized ossicles (hOss) have recently emerged as ectopic human bone/bone marrow microenvironments composed of MSCs and hematopoietic elements, enabling improved AML engraftment i*n vivo*. Nonetheless, those models typically exploit healthy MSCs largely because primary patient-derived cells exhibit diminished ossification potential. Here, we introduce OssiGel, a novel engineered human cartilage extracellular matrix produced by a mesenchymal cell line. We hypothesized that combining OssiGel with primary MSCs would enable the robust formation of hOss in healthy and malignant settings. We report that OssiGel supports fast and reproducible formation of hOss through endochondral ossification using MSCs isolated from both healthy and AML bone marrow samples. Within hOss, AML-MSCs were shown to persist long-term and to generate a human stromal microenvironment supporting the establishment of autologous AML hematopoietic cells, enabling the generation of autologous (autologOss). Single-cell RNA sequencing (scRNA-seq) revealed that autologOss recapitulate the majority of the AML blood populations observed in the corresponding patient bone marrow samples. Moreover, proof-of-concept sequencing of AML-MSCs from autologOss allowed predictive interrogation of interactions with leukemic populations and revealed the acquisition of an inflammatory-like phenotype, characteristic of AML. Taken together, our study establishes OssiGel as a novel platform for the robust engineering of autologous ossicles from AML patient samples. This provides a relevant tool to decipher patient-specific cellular interactions and drug responses, with potential translational value to broad hematological malignancies.

## Introduction

In healthy individuals, the bone marrow (BM) serves as a primary hematopoietic organ, producing and maintaining all blood cell lineages. It also provides a specialized niche that preserves hematopoietic stem cell (HSC) quiescence, self-renewal, and balanced differentiation. Within this niche, bone marrow mesenchymal stem/stromal cells (BM-MSCs) act as a key population giving rise to various lineages including osteoblasts, chondrocytes, adipocytes, perivascular and reticular stromal cells [1]. BM-MSCs hold key functions in healthy hematopoiesis by generating extracellular matrix (ECM) scaffolds that regulate HSC retention and proper presentation of pro-hematopoietic cytokines, and by directly interacting with HSCs to control their survival and differentiation [2].

During acute myeloid leukemia (AML) emergence, profound alterations of the BM microenvironment happen, progressively promoting leukemic growth to the detriment of their healthy hematopoietic counterparts [3,4]. Notably, BM-MSCs contributed to AML disease progression and therapy response by supporting leukemic cell survival, proliferation, and chemoresistance *via* cytokine networks, direct cell-to-cell contacts, and metabolic rewiring [5,6]. Transcriptomic and genomic profiling of human AML-derived MSCs revealed deregulated cytokine expression, adhesion molecules, and metabolic pathways [7]. Phenotypically, AML-MSCs also displayed altered differentiation potential and distinctive morphology compared with normal MSCs, suggesting niche remodeling by AML blasts, strongly supporting leukemia growth and therapy resistance [8,9]. Previous studies on human AML-MSCs have provided valuable snapshots of stromal alterations associated with the disease. However, most investigations remain cross-sectional, lacking longitudinal follow-up to capture the dynamic evolution of MSC reprogramming during disease. In addition, technical limitations include the low yield and variable quality of MSCs obtained from primary samples [10,11]. Human-relevant models that faithfully reproduce the leukemic BM niche remain scarce, particularly for myeloid malignancies.

*In vivo* investigations on these malignancies use xenograft transplantation into immunodeficient animals, as instrumental gold standard solution to study AML cell biology. However, in this setting, favorable and intermediate-risk patients often exhibit poor to no engraftment [12]. Most importantly, most models do not allow the functional study of the human niche upon disease development, which remain entirely murine. To address the limitations of conventional xenografts, engineered ectopic human BM niche models were proposed [13,14]. By reconstituting the human BM mesenchymal elements, they allow superior engraftment of hematological cancers including AML, myelofibrosis [15,16], multiple myeloma [17] and MDS [18]. Nevertheless, existing protocols to generate these humanized ossicles (hOss) are highly heterogenous, complex and lack reproducibility in forming mature bone and BM using patient BM-MSCs [19]. Notably, this limitation has so far challenged modeling the human leukemic BM niche changes using the hOss.

Recently, our group provided a standardized approach to generate hOss relying on a genetically modified human MSC cell line defined as Mesenchymal Sword of Damocles-Bone Morphogenetic Protein 2 (MSOD-B) [20]. MSOD-B hOss were shown to robustly support AML engraftment, correlated with the reconstitution of a complex human mesenchymal niche [21]. Moreover, we demonstrated that this system can be genetically engineered [22]. Despite the high degree of standardization conferred by the MSOD-B line, the immortalization of BM-MSCs may raise concerns on the biological relevance of the model and its capacity to fully reflect the patient-specific BM environmental changes. An attractive alternative proposes the formation of autologous humanized ossicles (autologOss), where both niche and leukemic cells originate from the same patient, thereby permitting advanced autologous niche interactions. So far, no reliable approach is available for the robust formation of autologOss. The main challenge lies in the required exploitation of AML patient BM-MSCs, which are notoriously difficult to expand *ex vivo* while maintaining their differentiation capacities [23]. To gain effectiveness in hOss formation, a promising strategy is based on exploiting the endochondral ossification [19]. During this process, MSCs undergo an intermediate cartilage state, then get gradually substituted by bone. However, the majority of MSCs from AML patients show an altered endochondral potential, thus failing to form hOss *in vivo* [24], or exhibit an impaired proliferation due to the frequent advanced age at diagnosis [10].

In this study, we aim to develop a new technology that enables the robust generation of hOss, bypassing the requirement for primary BM-MSCs to possess intrinsic chondrogenic potential. We hypothesize that embedding primary cells within a pre-established cartilage gel can restore their endochondral ossification capacity and their ability to form ectopic hOss. To this end, we introduce OssiGel as the first human-engineered cartilage hydrogel that, upon interaction with primary BM-MSCs, promotes consistent hOss formation via endochondral ossification and supports the establishment of a functional human mesenchymal niche.

In the context of AML, this strategy enabled the reproducible generation of autologOss, in which both the human MSCs and the engrafted AML cells originate from the same individual. These autologOss exhibited anticipated and new AML-MSC interactions, thus corroborating the translational value of the model, as shown using single-cell transcriptomic approaches. Taken together, we provide a robust platform for mechanistic investigations of human BM niche function in AML in a personalized manner.

## Results

### OssiGel is an engineered human cartilage hydrogel with robust osteoinductive properties

Conceptually, OssiGel was developed as a “revitalizing” hypertrophic cartilage hydrogel capable of restoring the chondrogenic-and thus endochondral ossification-potential of impaired patient-derived MSCs. To this end, MSOD-B cells were exploited as stable human MSC line with robust chondrogenic differentiation capacity [20]. MSOD-B were seeded on collagen scaffolds and *in vitro* primed for three weeks towards cartilage formation (**Figure 1A**). The resulting hypertrophic cartilage was subsequently lyophilized and cryo-milled to obtain a fine powder (**Figure 1B**), confirming the feasibility of transforming MSOD-B-derived cartilage into a processable raw material. Scanning electron microscopy analysis revealed that particles displayed a porous ECM microstructure and a size distribution falling within the 0.050-26.6 µm range (**Figure 1C**). Importantly, biochemical quantification demonstrated that despite freeze-drying and milling, the glycosaminoglycan (GAG) content was largely preserved during processing. In fact, comparable GAG levels were measured in lyophilized versus cartilage powder (11.1 ± 2.6 µg vs 10.4 ± 3.4 µg GAGs/mg ECM), suggesting that critical cartilage matrix components are retained in the final material (**Figure 1D**).

**Figure 1.**
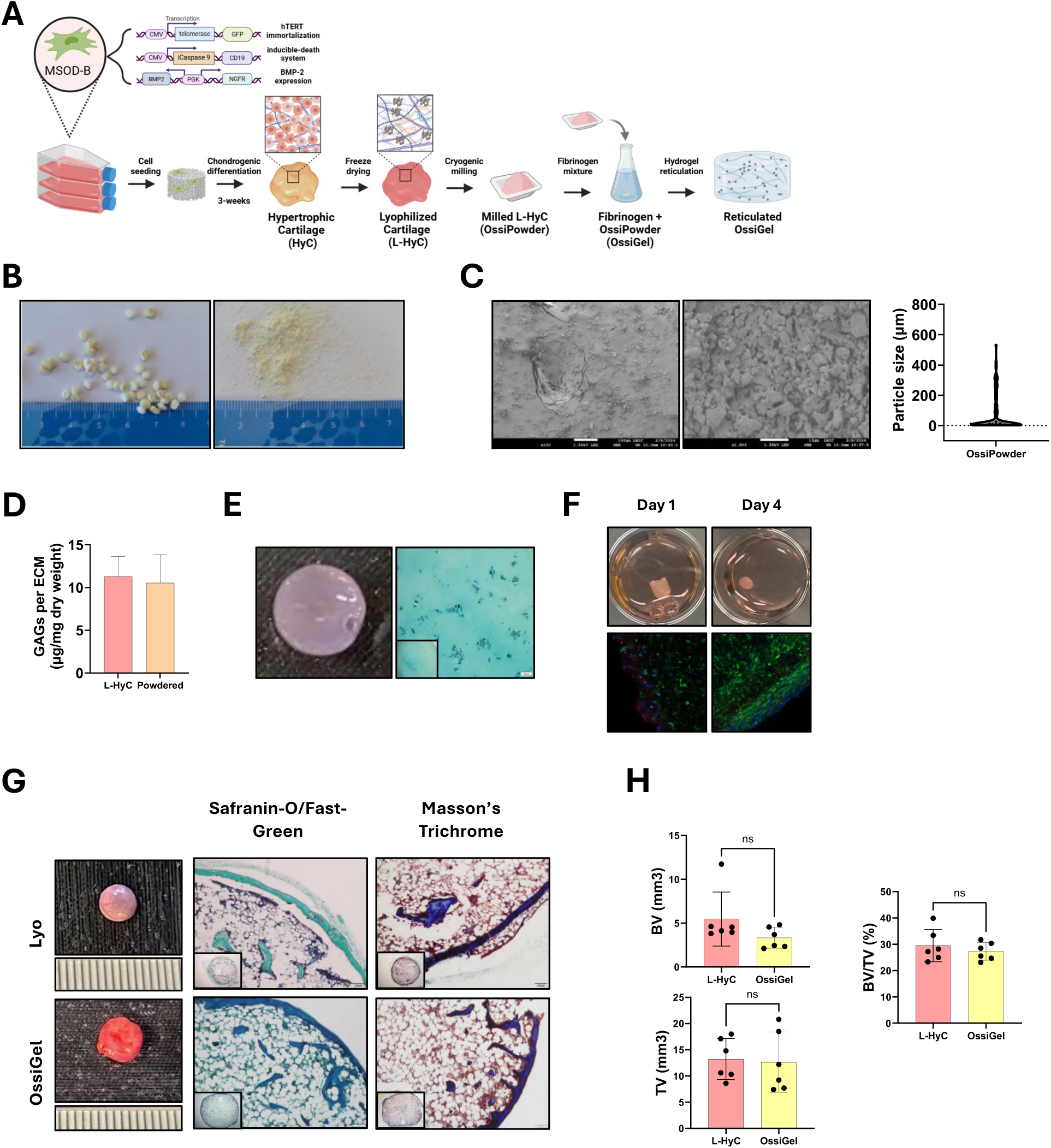
OssiGel is an engineered human cartilage hydrogel with robust osteoinductive properties. **A.** Schematic overview of the OssiGel generation process. **B.** Macroscopic view of the MSOD-B cartilage pellets after freeze-drying (left) and cryo-grinding (right). **C.** Scanning electron microscopy images of the cartilage powder and size distribution analysis. **D.** Glycosaminoglycan (GAG) quantification by lyophilized and powdered cartilage pellets, showing retention of main cartilage extracellular matrix (ECM) elements post-grinding. **E.** Macroscopic image and Safranin-O/Fast Green staining of powdered cartilage embedded in fibrinogen and thrombin post-reticulation. **F.** Macroscopic image and Live & dead staining of MSOD-B cells encapsulated within OssiGel and cultured *in vitro* for 1 and 4 days (*n*=4). **G.** Macroscopic appearance, Safranin-O/Fast-Green and Masson’s Trichrome staining of explanted tissues (*n*=6) from lyophilized pellets (Lyo) and OssiGel following ectopic implantation in immunodeficient mice for 6 weeks. **H.** Bone volume over total volume (BV/TV, %), bone volume (BV, mm^3^) and total volume (TV, mm^3^) of explanted tissues after 6 weeks of ectopic implantation in immunodeficient mice, assessed by micro-computed tomography (µCT) analysis (*n*=6).

Towards the generation of an injectable hydrogel material, we combined the powder with fibrinogen and demonstrate reticulation after thrombin addition, which led to a hydrogel with broadly distributed particles (**Figure 1E**). To assess its biocompatibility, we further embedded MSOD-B cells and cultured them in the reticulated hydrogel for up to 4 days. Over time, the encapsulated cells progressively condensed and proliferated, while live & dead staining did not indicate any cytotoxicity (**Figure 1F**). Together, these results demonstrate that engineered human cartilage can be transformed into a hydrogel by combination with fibrinogen and thrombin, which support the 3D culture of human cells.

The capacity of the hydrogel containing cartilage particles to drive endochondral ossification *in vivo* was further evaluated by ectopic implantation in BALB/C nude mice. As a control, lyophilized cartilage pellets (Lyo, not grinded) were also implanted. Six weeks post-implantation, both conditions generated hOss-like structures (**Figure 1G**). Histological analyses with Safranin-O/Fast-Green and Masson’s Trichrome stainings confirmed that implants underwent extensive remodeling leading to mature bone formation, as well as the establishment of a BM compartment. No cartilage remnants could be detected, while abundant trabecular bone and collagen-rich marrow-like areas were identified. Micro-computed tomography (µCT) quantification showed that both groups led to a similar bone formation (**Figure 1H**), with no statistical difference. The hydrogel group yielded a bone volume over total volume (BV/TV) fraction of 27.3 ± 3.3%, a bone volume (BV) of 3.33 ± 1.04 mm³, and a total volume (TV) of 12.6 ± 5.3 mm³, while those values reached 29.6 ± 6.1% (BV/TV), 5.80 ± 2.97mm³ (BV), 13.2 ± 3.8mm³ (TV) for the lyophilized. This indicates that the hydrogel containing cartilage particles retains sufficient instructive signals to drive endochondral ossification.

We further assessed the consistency of our approach by assessing various fibrin commercial providers, Sigma (S) and Baxter (B), at distinct concentrations (6 or 12 mg/mL). That resulted in four different formulations (S6: 6 mg/mL; S12: 12 mg/mL; B6: 6 mg/mL; B12: 12 mg/mL) which were combined with 2 mg of cartilage powder. Baxter fibrin, containing transglutaminase, yielded larger hydrogels with superior water retention (**Supp. Fig. 1A**). Consequently, more dispersed powder particles were observed within the fibrin network, as revealed by Safranin-O/Fast Green staining (**Supp. Fig. 1B**). Despite morphological differences, all formulations exhibited comparable mechanical properties (Young’s modulus; **Supp. Fig. 1C**). The osteo-inductivity of all formulations was further assessed *in vivo*. Following 6-week post-ectopic implantation in BALB/C nude mice, explants displayed similar morphology (**Supp. Fig. 1D**). Importantly, all of them achieved a complete remodeling into hOss, evidenced by Safranin-O/Fast-Green and Masson’s Trichrome stainings (**Supp. Fig. 1E**). Quantitative µCT confirmed an equivalent bone formation across all groups (**Supp. Fig. 1F**). Given this comparable biological performance, the 6 mg/mL Baxter fibrin formulation (formulation 3) was selected for all subsequent experiments. This selection was primarily based on qualitative macroscopic observations during surgery; the 6 mg/mL Baxter hydrogel demonstrated superior handling properties and better shape retention during ectopic implantation. Additionally, utilizing the lower fibrin concentration (6 mg/mL rather than 12 mg/mL) optimized material efficiency, allowing for a higher yield of constructs from a single reagent batch without compromising the regenerative outcome.

Collectively, these results support our central hypothesis that MSOD-B cartilage can be processed into a hydrogel, functioning as a novel biomaterial with intrinsic osteoinductive properties. We thus here defined the resulting material as OssiGel, an engineered human cartilage hydrogel priming endochondral ossification *in vivo*.

### OssiGel enables reproducible establishment of human-derived BM niches *in vivo*

OssiGel was evaluated for its capacity to generate human BM niches *in vivo*, in the form of hOss comprising BM-MSCs. To this end, healthy donor primary BM-MSCs at various amounts (0, 10^5^, 2×10^5^ and 10^6^ BM-MSCs) were encapsulated within OssiGel. Following reticulation, resulting scaffolds were implanted ectopically in BALB/C nude mice (**Figure 2A**). After 6 weeks, retrieved tissues were similar in size and morphology (**Figure 2B**), and consistent across BM-MSC donors (**Supp. Fig. 2A**). Safranin-O/Fast-Green staining indicated absence of remaining GAGs, suggesting a complete tissue remodeling (**Figure 2C**). This was confirmed using 3D µCT reconstruction, whereby all conditions led to mature hOss formation with cortical and trabecular bone structures (**Figure 2D**). Analysis of the trabecular structure indicated a trend for a BM-MSCs dose-dependent thickening of the trabecular structure in hOss (**Figure 2D**). Quantification of µCT (**Figure 2E**) revealed no significant effect of BM-MSC doses on the size of the hOss nor on the total amount of bone measured (BV/TV, No cells: 12.2 ± 4.5%; 100K: 17.0 ± 5.7%; 200K: 18.5 ± 4.1%; 1M: 17.8 ± 6.1%; ns). However, trabecular analysis confirmed the observed trend, with the 1M dose leading to distinctly thicker trabeculae (Trabecular thickness (Tb.Th): No cells: 157 µm; 100K: 158 µm; 200K: 225 µm; 1M: 286 µm; 1M vs others \*\*\*\**p*<0.0001) while trabecular spacing (Tb.Sp) was significantly elevated above 2×10^5^ cells when compared to acellular controls (\**p*<0.1; \*\**p*<0.01, **Figure 2F**). This suggests the active participation of human BM-MSCs in the trabecular deposition, as reported in previous model of endochondral ossification [25]. To assess the persistence of human BM-MSCs within the remodeled tissues, human nuclei (HuNu) staining was performed. HuNu-positive cells were detected in all constructs comprising human BM-MSCs (10^5^, 2×10^5^ and 10^6^ BM-MSCs), confirming the presence of implanted human cells *in vivo* (**Figure 2G**). Quantification of HuNu-positive cells showed a strong dependence of cell persistence on the initial BM-MSCs dose, with the highest number of HuNu-positive cells observed in the 10^6^ cell group (**Figure 2H**).

**Figure 2.**
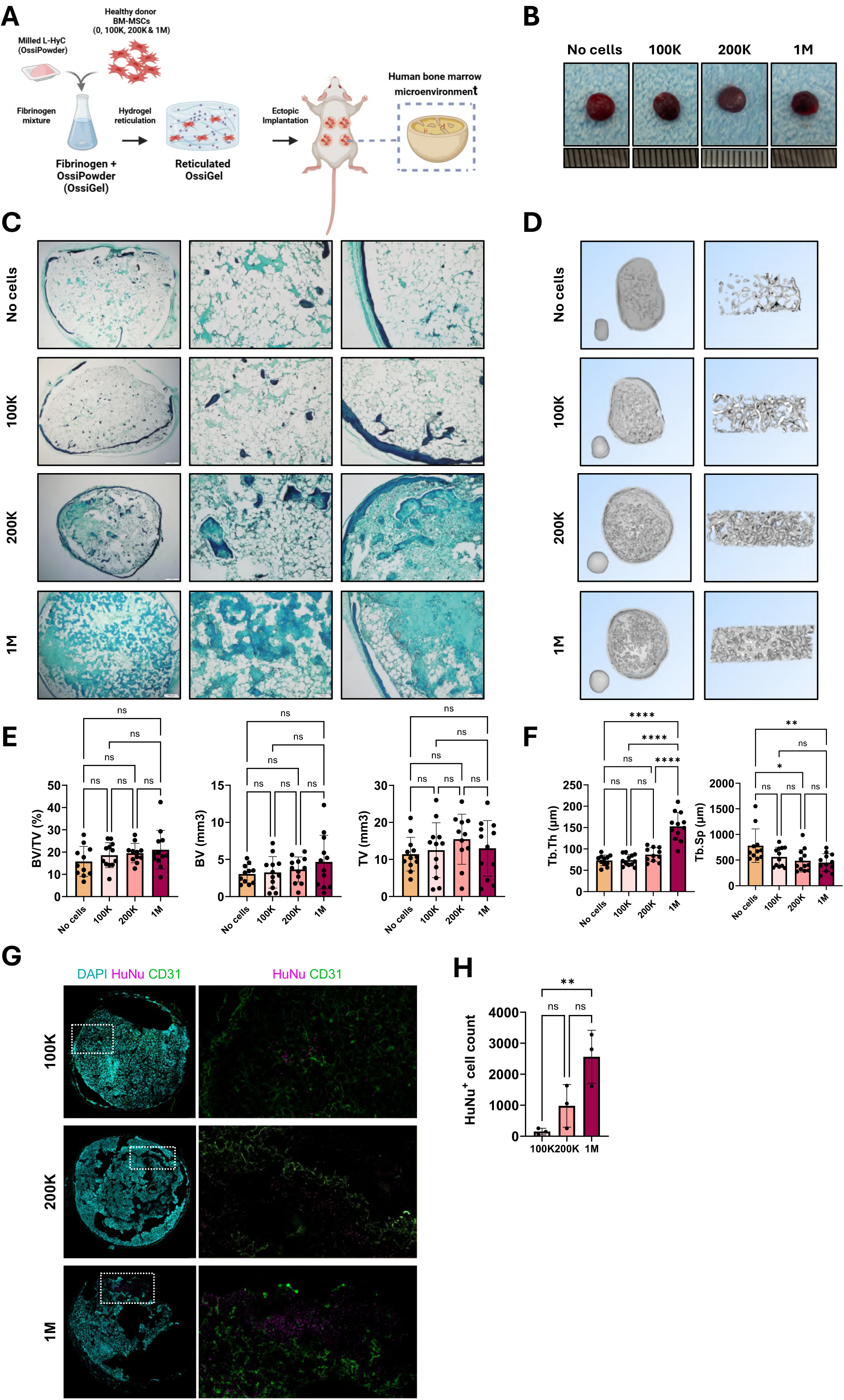
OssiGel enables the reproducible establishment of human-derived bone marrow niches within *in vivo* hOss. **A.** Schematic overview of the formation of hOss using OssiGel combined with various doses of human primary bone marrow mesenchymal stem/stromal cells (BM-MSCs). **B.** Macroscopic view of explanted tissues 6 weeks post-ectopic implantation in immunodeficient mice (*n*=4, 3 independent donors). **C.** Safranin-O/Fast-Green staining of explanted hOss resulting from different BM-MSCs doses. **D.** 3D reconstruction of full hOss using µCT (left) and of the trabecular region (right). **E.** BV/TV (%), BV (mm^3^) and TV (mm^3^) of explanted tissues after 6 weeks ectopic implantation in immunodeficient mice, assessed by µCT analysis (*n*=4, 3 independent donors). **F.** Trabecular thickness (Tb.Th) and trabecular separation (Tb.Sp) of explanted tissues 6 weeks post-ectopic implantation in immunodeficient mice, assessed by µCT analysis (*n*=4, 3 independent donors). **G.** Immunofluorescence staining of explanted hOss 6 weeks after ectopic implantation in immunodeficient mice formed with 1×10⁵, 2×10⁵ and 1×10⁶ BM-MSCs respectively. **H.** Quantification of Human Nuclei (HuNu) positive cell counts per 100 µm tissue section.

Another advantage of OssiGel lies in its potential injectability, offering the formation of ectopic human niches without the need for animal surgery. To evaluate this, we combined OssiGel with 1.5×10⁶ BM-MSCs using three different donors (D06, D09 and D13) and directly injected the mixture in the subcutaneous pocket of NSG mice (**Supp. Figure 2B**). After 6 weeks, tissues were successfully retrieved and exhibited round morphology but more heterogeneous shape, likely reflecting the variability from the injected area and possible spread until *in vivo* reticulation (**Supp. Figure 2C**). Remarkably, 3D µCT reconstruction confirmed complete hOss formation in all conditions with both cortical and trabecular structures (**Figure 2D**). Quantification of µCT confirmed a robust bone formation across donors (**Supp. Figure 2E**), with BV/TV, BV, and TV being comparable to hOss formed after implantation.

Taken together, these data validate the use of OssiGel for the formation of hOss, leading to the establishment of a human BM niche by implanted BM-MSCs in a dose-dependent manner, thus also influencing the trabecular composition. The ectopic niches are formed in only 6 weeks *in vivo* and can be obtained by direct injection of the OssiGel/BM-MSCs mixture.

### OssiGel allows consistent generation of AML hOss

The formation of hOss using primary BM-MSCs from AML patients (AML-MSCs) is challenging, due to their altered chondrogenic potential. To this end, we investigated the potential of OssiGel in driving endochondral ossification of AML-MSCs, towards the generation of patient hOss. AML-MSCs from 9 patients with diverse cytogenetic backgrounds (**Table 1**) were isolated, *in vitro* expanded and mixed with OssiGel (**Figure 3B**) prior to subcutaneous implantation into immunodeficient NSG mice. In this study, we also interchangeably used lyophilized cartilage pellets (Lyo), loaded with AML-MSCs and subsequently implanted (**Supp. Fig. 3A**). After 6 weeks *in vivo*, the CD3^-^CD45^+^ blood fraction from the same AML patient was injected directly into the hOss, in order to reconstitute the patient-specific BM microenvironment in an autologous fashion (**Figure 3A**). On average, each tissue comprised 1×10^6^ AML-MSCs, ranging from 2.5×10^5^ to 1.36×10^6^ cells (**Table 2**).

**Figure 3.**
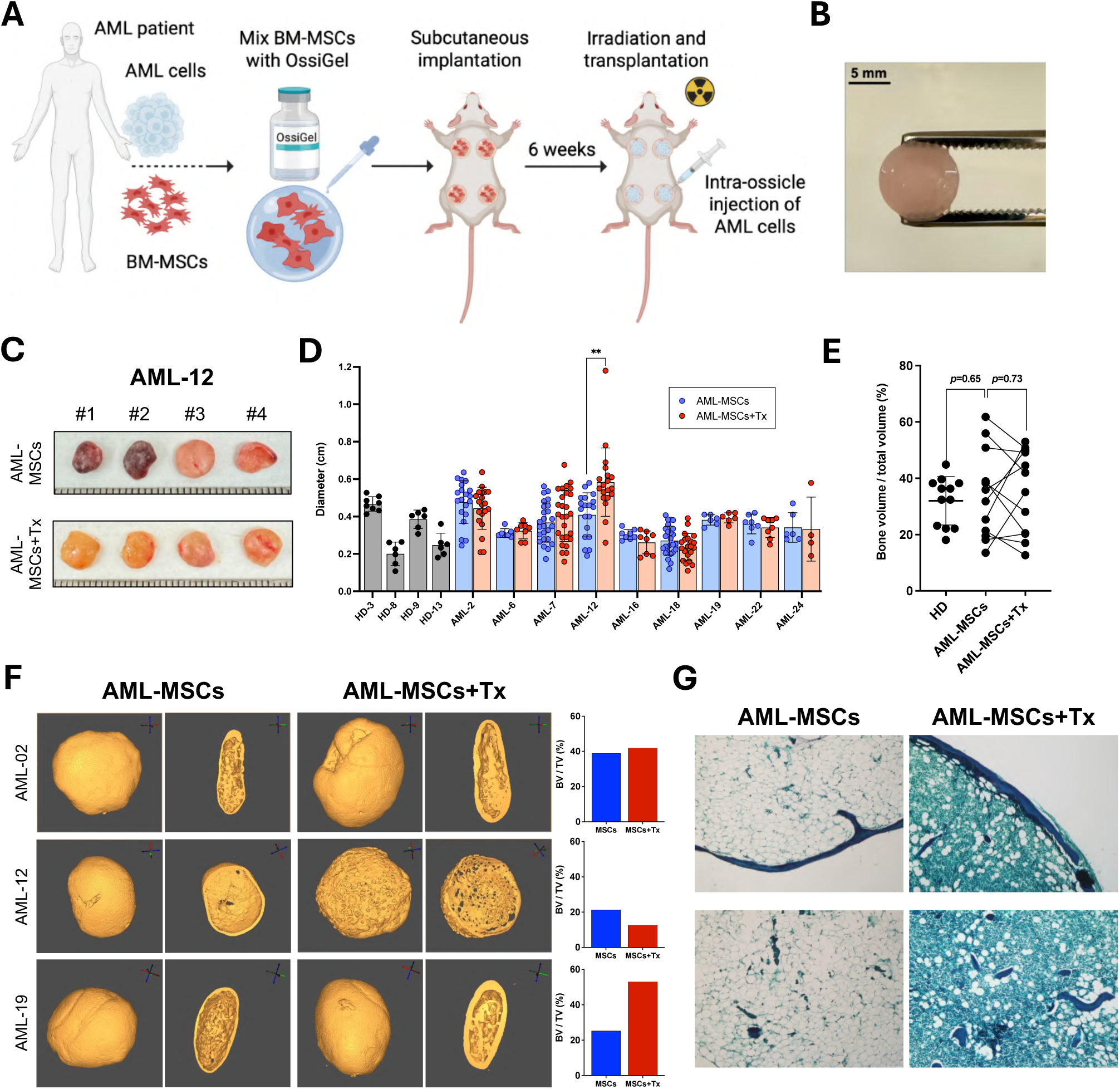
Robust and reproducible generation of acute myeloid leukemia (AML) patient-derived hOss. **A.** Experimental scheme for the generation of hOss from AML patient bone marrow samples. **B.** Macroscopic picture of an OssiGel containing AML-MSCs. **C.** Representative macroscopic images of hOss with and without AML cell transplantation 12 weeks after explantation. **D.** Diameter of hOss from healthy donors (*n*=3) and patients with and without AML (*n*=9) cell transplantation. Each dot represents one hOss (*n*=4-20 hOss per condition). \*\**p*<0.01. **E.** Quantification of BV/TV of hOss from healthy donors (*n*=3) and AML patients (*n*=5) with and without AML cell transplantation. Each dot represents one hOss. **F.** µCT images of hOss from 3 different patients with and without AML cell transplantation after explantation and quantification of the BV/TV ratio. **G.** Representative images of Safranin-O/Fast Green staining on paraffin sections from hOss. Top: peripheral area, bottom: central area of the hOss.

**Table 1.** Healthy donors and AML patients’ information.

|  | Age | Gender | Cytogenetics | ELN 2022 risk stratification | Blasts in BM |
| --- | --- | --- | --- | --- | --- |
| HD-03 | 28 | Female |  |  |  |
| HD-08 | 29 | Female |  |  |  |
| HD-09 | 19 | Male |  |  |  |
| HD-13 | 23 | Female |  |  |  |
| AML-2 | 39 | Female | normal karyotype, mutations in RUNX1 (VAF 95%), EZH2 (6%), EZH2(9%) , EZH2 (10%), IDH2 (46%), PIK3CD (5%), TRRAP (3%) | Adverse | 75% |
| AML-6 | 73 | Female | 42-44, XX, -4, -5, del(6)(q-?), -8, -12, -13, -15, -17, +3-4 mar [cp19]/46, XX[6], NGS panel: TP53 (67%), DNMT3A (36%), IDH1 (40%), STAG2 (15%) | Adverse | 78% |
| AML-7 | 75 | Female | 45-50,X,-X, idic(X)(q13),+8,+11,+13, +19[cp24]/46,XX[1] ; NGS: ASXL1 (4%), NRAS (27%), TET2 (47%), TET2 (48%), SRSF2 (46%) | Adverse | 51% |
| AML-12 | 59 | Male | 46, XY [25]; FLT3-ITD mutated with allelic ratio 1,73 (high), RUNX1 (42%), DNMT3A (45%), TET2 (45%) | Adverse | 74% |
| AML-16 | 70 | Female | inv (3)(q21;q26)/t(3;3)(q21;q26); FISH positive for MECOM-gene | Adverse | 14% |
| AML-18 | 59 | Male | 48, XY, +4, del (9)(q12;q22), +21;[25]; AML, NPM1 (41%), TET2 (32%), TET2 (46%), FLT3-ITD mutated with allelic ratio of 0,67 (high) | Intermediate | 90% |
| AML-19 | 69 | Male | 46, XY, [25]; NGS: RUNX1 (74%), ASXL1 (37%), BCORL1 (36%), BCORL! (32%), CSF3R (12%), ZBTB (79%), SAMD9L (35%) | Adverse | 53% |
| AML-22 | 77 | Male | complex karyotype with del (5), del 17, del 18 ; NGS: TP53 (61%), TET2 (42%), ETV6 (43%) | Adverse | 32% |
| AML-24 | 46 | Male | 46, XY, inv (16), (p13; q22)/46, XY; CBFB/MYH11 AML | Favorable | 35% |

**Table 2.** Number of MSCs, number of AML cells transplanted, time of explantation and number of hOss retrieved from each experiment.

|  | Scaffold type | Nb of MSCs in each construct (millions) | Nb of AML cells in each hOss (millions) | Explantation (weeks after transplantation) | % of hOss retrieved |
| --- | --- | --- | --- | --- | --- |
| HD-03 | Lyophilised | 0.25 / 0.5 |  | 4 weeks after implantation | 100 |
| HD-08 | OssiGel | 2.7 |  | 8 weeks after implantation | 100 |
| HD-09 | OssiGel | 2 |  | 8 weeks after implantation | 100 |
| HD-13 | OssiGel | 2.7 |  | 8 weeks after implantation | 100 |
| AML-2 | OssiGel | 1 | 2.94 | 29 | 100 |
|  | Lyophilised | 0.5 | 1 | 17 | 100 |
| AML-6 | Lyophilised | 0.253 | < 0.01 | 33 | 93.8 |
| AML-7 | OssiGel | 1 | 1.1 | 27 | 87.5 |
| AML-12 | OssiGel | 0.62 | 1.88 | 12 | 100 |
|  | Lyophilised | 1.12 | 4.95 | 18 | 87.5 |
| AML-16 | Lyophilised | 0.7 | 0.2 | 26 | 100 |
| AML-18 | OssiGel | 0.6 | Intravenous (10.41/mouse) | 32 | 85.6 |
|  | Lyophilised | 1.36 | 4.2 | 21 | 100 |
| AML-19 | Lyophilised | 0.58 | 1.723 | 19 | 100 |
| AML-24 | OssiGel | 0.25 / 0.5 | 1 | 16 | 62.5 |
| AML-22 | Lyophilised | 0.417 | 2 | 32 | 93.8 |

Remarkably, after 4-6 weeks *in vivo*, AML-MSCs generated patient-derived hOss across all AML cases, with an ossification success rate of 93.12 ± 10.75%. This represents a total of 269 patients hOss retrieved at explantation (**Table 2**). Macroscopic observations reveal a rather homogenous morphology among replicates from the same patient. However, a consistently reduced red blood cell fraction was macroscopically evident in hOss transplanted with AML (AML-MSCs+Tx), compared to non-transplanted hOss (AML-MSCs) (**Figure 3C, Supp. Fig. 3B**). Measurements of hOss diameters indicated a similar size across patient hOss including those generated using healthy BM-MSCs (HD) (**Figure 3D**). Of note the transplantation of AML blood cells did not impact the diameter of resulting hOss, with the exception of one patient (AML-12) that displayed a significant size increase in transplanted tissues.

Quantitative µCT analysis (**Figure 3E**) confirmed no significant difference in hOss dimensions (BV/TV) between healthy donor-derived and AML hOss (32.0 ±8.6% vs 34.3 ±15.7%), nor between non-transplanted (AML-MSCs) and transplanted AML hOss (AML-MSCs+Tx) (34.3 ±15.7% vs 36.5 ±14.9%). The 3D reconstruction of patient-derived hOss (**Figure 3F**) confirmed the formation of mature bone tissue but also revealed distinct, patient-specific effects of transplantation on the overall bone architecture. While some hOss appeared unaffected (AML-02), AML transplantation led to either increased bone content (i.e. AML-19) or reduced BV/TV accompanied by osteolytic lesions (i.e. AML-12, **Figure 3F**). Histological analyses with Safranin-O/Fast Green confirmed the formation of mature hOss comprising a BM cavity which seem to be infiltrated by transplanted AML cells (**Figure 3G**).

Overall, these data demonstrate the feasibility of generating AML patient hOss in a reproducible fashion using OssiGel. While the protocol is robust, these hOss exhibited patient-specific features, illustrated by varying sizes and distinct patterns of bone formation among patients.

### Robust AML cell establishment and long-term persistence of AML-MSCs is evident in autologous AML niches

Previous hOss models demonstrated superior engraftment of AML patient cells as compared to standard mouse bones [15,16,26]. However, none of those exploited AML-MSCs for ossification, thus failed at recapitulating the patient-specific BM niche. Following bone formation in hOss using AML-MSCs, we next assessed whether these MSCs persisted within the established tissue, and whether AML blood cells successfully engrafted and circulated in other hematopoietic organs of the host.

Following successful bone formation confirmed by *in vivo* µCT, each hOss was transplanted with their corresponding AML patient, on average with 1.93×10^6^ hematopoietic cells (ranging from 2×10^5^ to 4.95×10^6^, **Table 2**). Engraftment was monitored in peripheral blood and/or in hOss (through hCD45 expression), and upon disease detection, hOss and corresponding mouse femurs were collected and processed for subsequent analyses (between 12 and 33 weeks post-transplantation, **Figure 4A**). Human blood engraftment was typically detected at low level in peripheral blood; only two out of nine AML patients displayed more than 0.1% of hCD45^+^ cells out of total hematopoietic cells at the time of explantation (**Figure 4B**). However, seven out of nine AML patients showed long-term establishment of human hematopoietic cells in hOss at explantation, displaying more than 1% of human CD45^+^ cells. Strikingly, the percentage of hCD45^+^ cells was significantly higher in patient hOss compared to the corresponding mouse femurs across all experiments (Wilcoxon matched-pairs signed-rank test, W=36, *n*=8, *p*=0.0078, **Figure 4C, Supp. Fig. 4A**).

**Figure 4.**
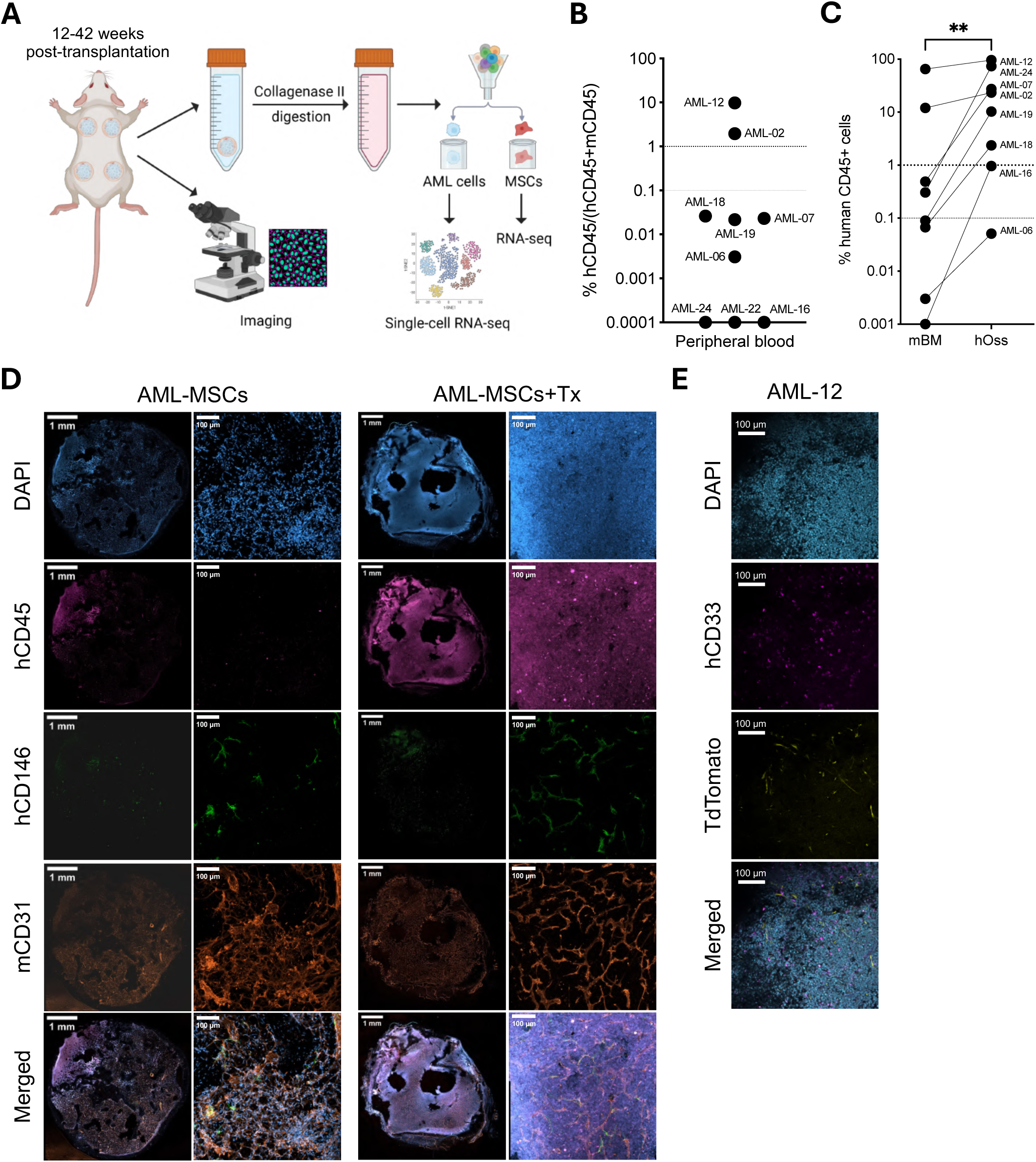
Robust AML cell establishment and long-term persistence of AML-MSCs is evident in autologous AML niches. **A.** Experimental scheme for the processing of hOss from AML patient at endpoint of experiments (between 12 and 33 weeks after transplantation). RNA-seq: RNA-sequencing. **B.** Percentage of engraftment in peripheral blood of NSG mice bearing hOss with AML transplantation (autologOss, for autologous ossicles) at endpoint of experiments. Each dot represents one AML patient; *n*=2 to 5 mice per patient sample. **C.** Percentage of hCD45^+^ cells in autologOss versus corresponding mouse femurs at endpoint of experiments. Each dot represents one AML patient; *n*=2 to 5 mice per patient sample. **D.** Representative confocal images of hCD45^+^ cells and CD146^+^ MSCs within hOss with and without AML cells 12 weeks after transplantation. Nuclei are stained with DAPI. Maximum intensity projection of a Z-stack at magnification x10 (whole sections) or x20 (details). **E.** Representative confocal images of hCD33^+^ cells and TdTomato^+^ MSCs within autologOss. Single stack images at magnification x10.

The low levels of circulating AML blood cells, despite successful engraftment, suggest that these cells are largely confined within the established hOss where supporting AML-MSCs persist. This was confirmed by immunofluorescence analyses of patient hOss sections, revealing the presence of human stroma expressing the hCD146 marker (**Figure 4D, Supp. Fig. 4B**). These AML-MSCs were detected in both non-transplanted and transplanted patient hOss, and as anticipated, were absent from their respective mouse femurs (**Figure 4D, Supp. Fig. 4B**). Beyond AML-MSCs, immunofluorescence also evidenced a dense vascular network of murine origin (mCD31) of similar density in hOss and murine native bones. Transplanted hOss exhibited an altered BM, crowded with human blood cells (hCD45), typical of the profound alterations in AML BM. To confirm long-term persistence of AML-MSCs in patient hOss beyond the sole hCD146 expression, we transduced AML-MSCs (7 patients) with a lentiviral vector encoding TdTomato prior to hOss generation (**Supp. Fig. 4C**). After tissue explantation, immunostainings revealed the persistence of TdTomato^+^ AML-MSCs cells for up to 26 weeks post-transplantation (**Figure 4E, Supp. Fig. 4D**).

Taken together, these findings demonstrate the successful generation of patient-derived hOss in which both the hematopoietic and mesenchymal compartments are robustly reconstituted. This establishes an autologous ossicle (autologOss) model that supports long-term AML maintenance and provides a powerful platform to functionally interrogate AML/MSC interactions.

### Single-cell transcriptomic analysis of AML blood cells from autologOss reveals broad blood lineage reconstitution

Next, we wanted to assess patterns of AML blood reconstitution using single-cell RNA transcriptomic (scRNA-seq) tools, following fluorescence-activated cell sorting for hCD45^+^ (hematopoietic fraction) or mCD45^-^hCD45^-^TdTomato^+^ for MSC-derived cells (**Figure 4A, Supp. Fig. 4E**). Hematopoietic CD45^+^ cells were sorted from the originally collected AML patient BM aspirates, as well as using BM isolated from both autologOss and corresponding mouse femurs. The stromal TdTomato^+^ fraction was sorted from autlogOss as well as *in vitro* expanded MSCs from the same patient. Patient data from two independent experiments using the same AML patient sample that showed substantial engraftment both in autologOss and mouse bones (AML-12) are shown in **Figure 5**.

**Figure 5.**
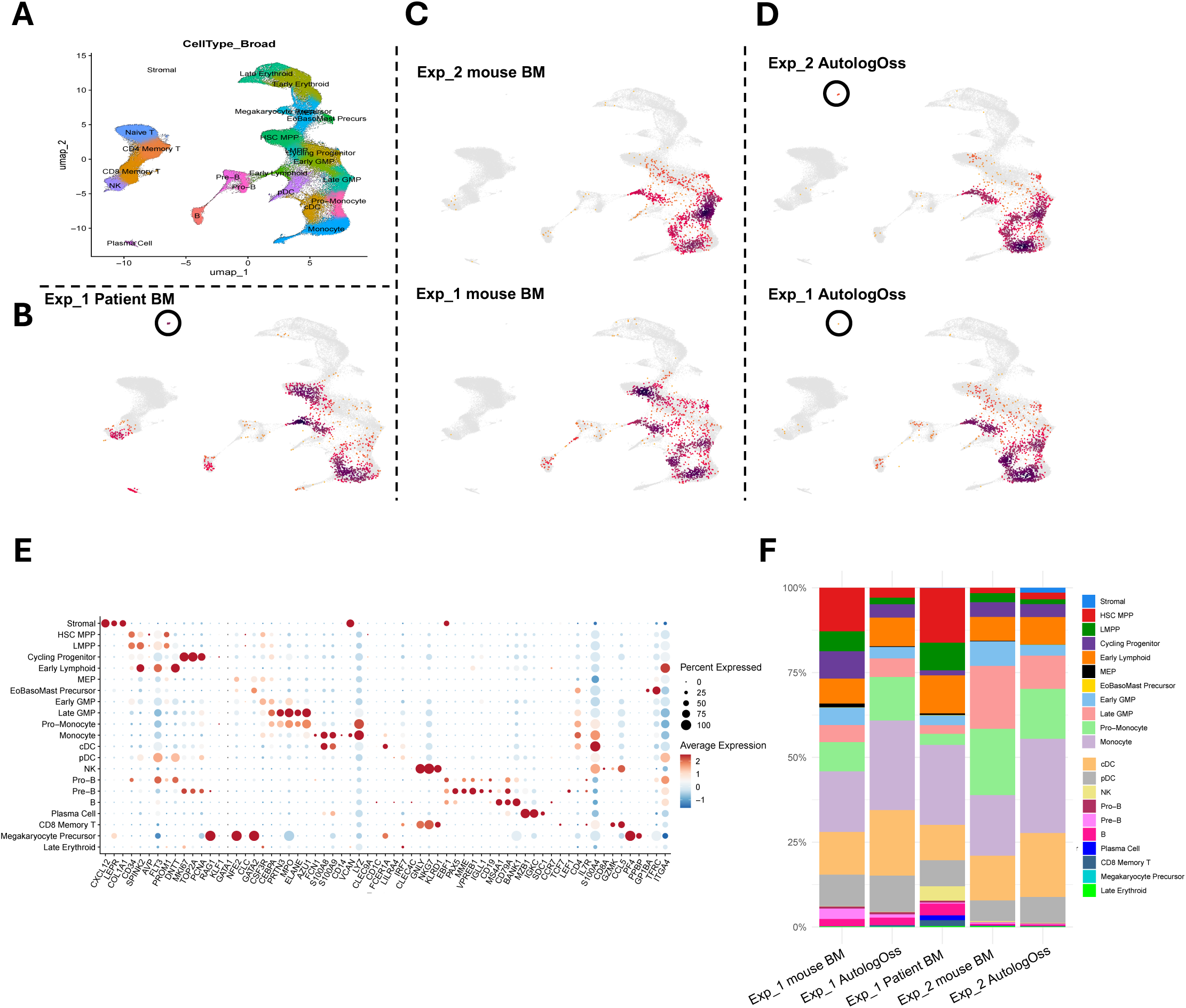
Single cell transcriptomic analysis of AML blood cells from autologOss reveals broad blood lineage reconstitution. **A.** Uniform Manifold Approximation and Projection (UMAP) of the reference dataset (Zeng *et al*., Blood Cancer Discov, 2025), colored by annotated cell type. **B-D.** UMAP density maps of the reference UMAP space showing the distribution of cells from each sample: Patient bone marrow (B), hCD45^+^ cells isolated from murine femurs from two independent experiments (C, top and bottom), and hCD45^+^ cells cells isolated from autologOss from two independent experiments (D, top and bottom). Circles in B and D highlight the presence of stromal cells. **E.** Dot plot displaying scaled expression of key marker genes across annotated cell populations. Dot size reflects the proportion of cells expressing each gene; color intensity indicates mean scaled expression. **F.** Stacked bar plot showing the proportional composition of annotated cell types across individual samples, with each color representing a distinct cell population.

Following per-sample quality control and dataset integration, the merged dataset comprised a total of 21,063 human cells. To assign cell type identities, we mapped our data onto a publicly available BM reference atlas [27] (**Figure 5A**), which provides a comprehensive representation of BM cell types. Density mapping onto the reference revealed a consistent myeloid bias across patient BM (**Figure 5B**), mouse femurs (**Figure 5C**), and autologOss (**Figure 5D**). Density maps also revealed a high reproducibility across the two experimental repeats, with similar reconstitution of various blood populations. Importantly, a rare human MSC cluster was detected exclusively in autologOss and in the original patient material, consistent with the anticipated presence of a human BM niche in engineered tissues niche (**Figure 5B-D, Supp. Fig. 5A**). To validate predicted cell type identities, we further performed differential gene expression analysis across identified clusters (**Supp. Fig. 5C**) and assessed the expression of canonical lineage marker genes for each population (**Figure 5E**). Lastly, a cell type distribution analysis across samples allowed evaluating potential differences between patient BM, mouse BM and autologOss (**Figure 5F, Supp. Fig. 5B**). This revealed that the most pronounced compositional differences consisted in the pro-monocyte and monocyte compartments, which showed higher representation in the hOss. Femur engraftment displayed greater inter-experimental variability, particularly at the HSC-MPP and LMPP populations. Nevertheless, both niches failed to recapitulate other cell types present in the patient BM such as T-cells, as CD3^+^ cells were depleted prior to transplantation. Additionally, B-cell populations also showed a high degree of inter-experimental variation.

Overall, scRNA-seq confirmed the reconstitution of AML BM cellular clusters in both mouse femurs and autologOss. Cluster compositional differences between samples existed but remained homogenous across experimental repeats. Despite the fact that murine femur and autologOss niches were able to successfully host AML, autologOss may provide a standardized platform for AML modeling with composition similarities to the patient’s original BM. This validates the possibility to exploit autologOss as a tool to decipher AML BM function at single-cell resolution.

### AutologOss offer the personalized study of MSC / AML blood cell interactions

The detected MSC population in the scRNA-seq dataset was further characterized, together with patient MSCs *in vitro* expanded post-isolation from the patient BM aspirates. We could thus uncover transcriptional changes that occurred in MSCs following *in vivo* implantation and AML engraftment. The MSC cluster from the complete dataset was isolated and a sub-clustering analysis was performed. The resulting MSC sub-dataset, comprising 967 cells, resolved into three distinct clusters: two composed of *in vitro*-expanded MSCs and one composed of MSCs isolated from autologOss (**Figure 6A**). Cell identity of autologOss was confirmed by the expression of a panel of canonical stromal marker genes (**Figure 6B**), including *PDGFRB, CXCL12, ENG* and *THY1*, and the lack of hematopoietic cell markers.

**Figure 6.**
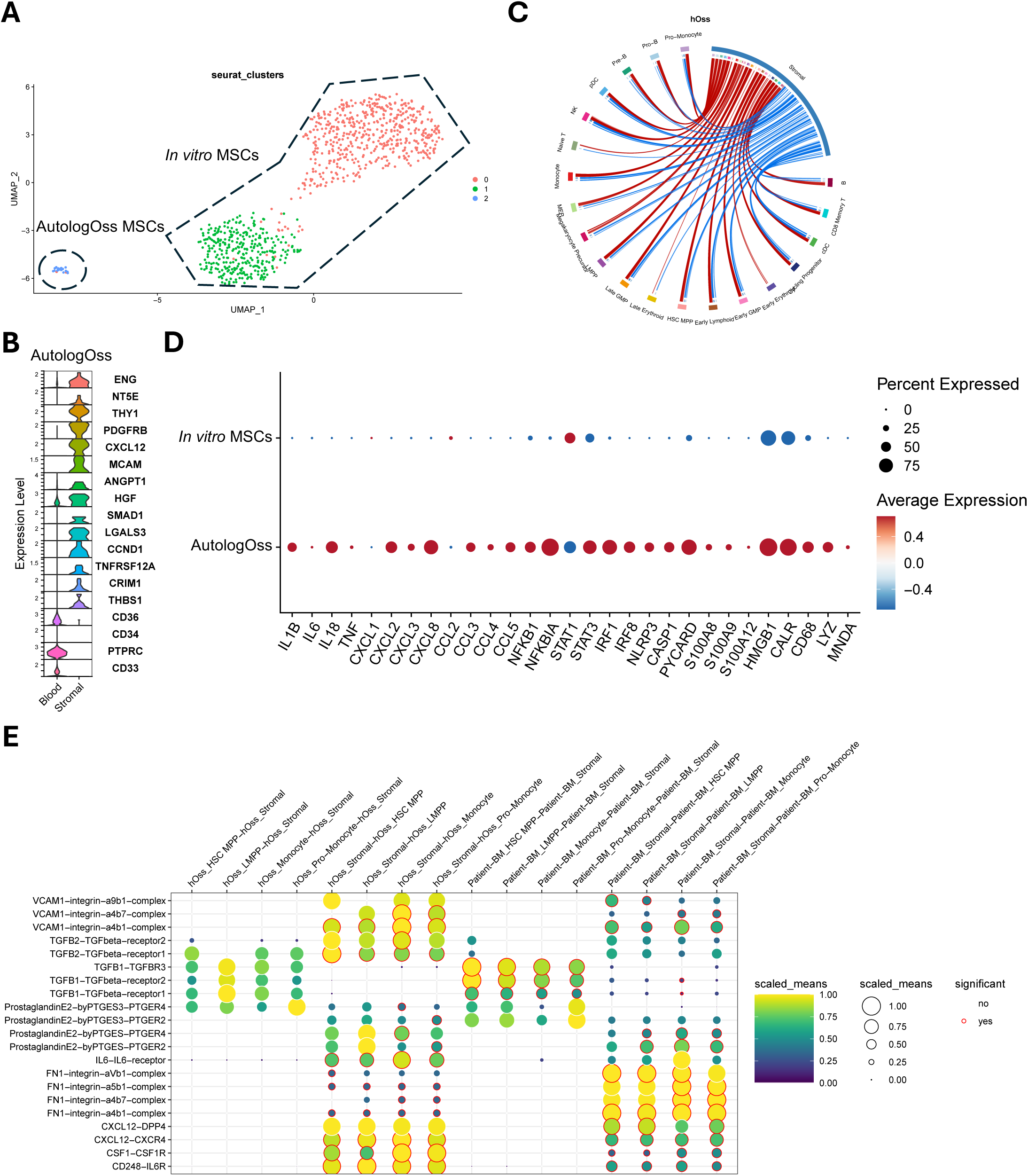
AutologOss offer the personalized study of MSC / AML blood cell interactions. **A.** UMAP embedding of stromal cells isolated from the broader single-cell dataset, colored by transcriptional subpopulation. **B.** Violin plot of key stromal and blood markers on celltypes isolated from autologOss, Blood represent all autologOss bone marrow-derived cell types. **C.** Circos plot illustrating the predicted intercellular communication network celltypes present in the autologOss and the respective stromal cells **D.** Dot plot depicting scaled expression of key inflammatory mediators in MSCs derived from expanded MSCs from patient BM (top) and stroma isolated from autologOss (bottom). **E.** Dot plot of enriched cell-cell communication pathways between MSCs and selected blood populations, with dot size indicating interaction probability and color reflecting mean expression of the corresponding ligand-receptor pairs.

Furthermore, differential gene expression analysis across clusters confirmed that autologOss-MSCs primarily consist in a stromal population with increased *CXCL12* and key ECM gene expression (*COL1A1, COL3A1, VCAM1, THY1*, **Supp. Fig. 6A**). Of note, the MSC population in autologOss acquired a pro-inflammatory profile, characterized by the expression of a wide range of proinflammatory genes (including *IL1B, IL18, NFKB1, NFKBIA, STAT*, **Figure 6D**), commonly upregulated in JAK-STAT and NF-κΒ activated states and as previously reported in AML [28]. In sharp contrast, *in vitro* expanded patient MSCs did not exhibit such profile, confirming loss of inflammatory profile upon *in vitro* culture. In parallel, blood populations established in autologOss similarly displayed a significantly higher expression of proinflammatory genes, as compared to the patient BM and mouse femurs (**Supp. Fig. 6B**).

We next applied CellphoneDB [29] to predict cell-to-cell communication on our dataset on a per-organ basis. The previously performed differential gene expression analysis served as input for selection of ligand-receptor pairs of stroma and blood populations. We observed that MSCs formed an interactive hub, interacting the most and broadly with all human hematopoietic populations (**Figure 6C, Supp. Fig. 6C–D**). Interestingly, this pattern was conserved in *in vitro* expanded AML-MSCs (**Supp. Fig. 6C**). Previously reported axes of communication were identified and preserved in both *in vitro-* and autologOss-derived MSCs, including but not limited to the CXCL12-CXCR4 pathway, the key anchor pathway linking MSCs and hematopoietic cells. Interestingly, signaling pathways led by the blood compartment remained mostly consistent across conditions, showing high expression of the TGFB1-and prostaglandin E2-related genes. Interactions led by MSCs showed sharp differences, namely autologOss-MSCs displayed an upregulation of VCAM1-integrin, IL6-IL6R and TGFB2-TGFBR2 complexes, which was not observed with *in vitro* expanded MSCs, indicating potential AML-MSC specific interactions in autologOss. Finally, we observed a predicted increased expression of the CSF1-CSF1R complex between autologOss-derived stroma and monocyte/pro-monocyte populations; in line with our reported higher content of monocyte/pro-monocyte cells in autologOss (**Figure 6E, Supp. Fig. 6E**).

Overall, MSCs within the autologOss niche display a distinct *in vivo*-induced transcriptional state characterized by a pro-inflammatory and ECM-enriched profile that is not preserved after *in vitro* expansion. These MSCs act as key communication hubs, exhibiting enhanced and disease-associated ligand-receptor interactions with hematopoietic cells, including AML-relevant signaling axes. Together, these results highlight a context-dependent remodeling of the stromal compartment that may contribute to shaping the leukemic microenvironment.

## Discussion

Collectively, our findings demonstrate that OssiGel can act as a human ECM to prime endochondral ossification of BM-MSCs. The technology was demonstrated to enable the formation of autologOss, recapitulating both the hematopoietic and mesenchymal niche features of AML patients. Single-cell transcriptomics of autologOss demonstrated recapitulation of MSC-leukemic cells, as well as the acquisition of an inflammatory phenotype by MSCs, validating the possibility to exploit autologOss as a tool to decipher AML BM function at single-cell resolution and in a personalized fashion.

Endochondral ossification is the developmental process through which MSCs form and regenerate bone. Although this capacity is intrinsic to BM-MSCs, it critically relies on their capacity to undergo chondrogenic differentiation. In fact, a substantial proportion of primary MSCs from healthy donors fail to form cartilage and thus to efficiently undergo endochondral ossification *in vivo* [14]. This variability represents a major limitation for the reliable generation of human BM hOss.

In this study, we introduce a fundamentally novel concept to generate hOss using a highly standardized endochondral-priming hydrogel, derived from the engineered human MSC line MSOD-B [20]. Because it originates from a stable cell line, this provides an unlimited and consistent source for cartilage formation subsequently transformed into a gel-like biomaterial. Importantly, we showed that the conversion into a hydrogel preserves its endochondral capacity, enabling embedded primary MSCs to be effectively primed toward bone formation and niche establishment. A core advantage of the resulting OssiGel lies in its injectability, which eliminates the need for animal surgery while ensuring a high degree of reproducibility. Using OssiGel, we demonstrate the formation of a human BM niche *in vivo* within 4-6 weeks. Moreover, the system is tunable: varying the proportion of BM-MSCs allows dose-dependent modulation of hOss formation and trabecular architecture. As a comparison, existing ossicle-generation protocols can require up to 14 weeks to achieve ossification [14], making our method both substantially faster and highly reproducible.

Because of the primary MSCs variability, existing hOss models rely on selected batches of healthy BM-MSCs to establish bone and marrow tissues [14,19]. In fact, MSCs from aged or leukemic patients are further compromised, with impaired functional reprogramming, mainly characterized by diminished osteogenic differentiation and a concomitant increase in adipogenic lineage [9,10,24,30]. This adds additional complexity to experimental design and highly impairs reproducibility in existing studies. Our lab partially addressed this limitation by using MSOD-B as a stable, standardized cell source for hOss formation [21]. However, this did not overcome the challenge of reconstructing a patient-specific BM niche. Here, we demonstrated that OssiGel restored the ossification potential of AML-MSCs, leading to the first reliable approach to generate autologOss, defined by reconstitution of both the mesenchymal niche and hematopoiesis from the same patient. This overcomes a key barrier and establishes a versatile and standardized platform for studying human BM in a patient-specific context. Notably, diameter and BV/TV measurements of AML-derived hOss revealed substantial interpatient variability, indicating heterogeneity in structural bone formation across samples. In our experimental set, the presence of MSCs appeared to influence both overall bone integrity and the spatial distribution of marrow compartments within the ossicles, suggesting a functional contribution of MSC content to tissue organization. However, the mechanisms underlying this effect, as well as its biological determinants and impact on niche fidelity, remain to be elucidated in greater depth.

In terms of AML engraftment, we reported consistent disease establishment following intra-ossicle delivery of AML hematopoietic cells. This approach overcomes a key limitation of conventional xenotransplantation systems, in which many primary AML samples engraft poorly or inconsistently [31]. Compared with the MSOD-B hOss model, we observed similar engraftment efficiency and AML burden within the hOss BM [21], and we further compare AML populations detected in hOss and murine femoral BM compartment. However, these observations should be interpreted with consideration of the current experimental design, as femoral AML populations may in part reflect cells circulating from the ossicles, where the disease was implemented *in vivo*. Future studies incorporating unbiased intravenous delivery approaches will help to more definitively assess relative engraftment across anatomical sites.

Despite the advantages conferred by the autologOss system, a subset of patient samples still failed to establish disease. This variability reflects the marked heterogeneity of AML and indicates that successful engraftment is not determined solely by the microenvironment. Indeed, AML progression *in vivo* also depends on leukemia-intrinsic factors, including cytogenetic features and disease status at diagnosis, such as the proportion of blasts present in patient samples at collection [12,31]. Of note, OssiGel-derived hOss from AML patients were successfully generated irrespective of the patient’s ELN risk classification. Another contributing factor for disease establishment may be the quantity of patient-derived cells introduced into the system. We reported a dose-dependent persistence of BM-MSCs within hOss, suggesting the benefit of implanting high amount of MSCs. However, the approach remains constrained by the limited volume of patient biopsies and the variable efficiency of AML-MSC isolation and expansion *in vitro*. Similarly, increasing the transplanted hematopoietic fraction could further enhance or accelerate engraftment, but the reliance on the availability and quality of patient-derived material persists.

Nonetheless, the OssiGel model enables the engineering of an autologous BM niche and subsequent single-cell profiling of both hematopoietic and MSC compartments. Comparison with the original patient BM sample demonstrated reliable reconstitution of AML blood in autologOss, but also in mouse BM. Minor differences were observed, in particular a higher pro-monocyte/blast content in autologOss, suggesting an impact of the human niche on disease reconstitution *in vivo*. Assessing additional patient samples would allow identifying clear patterns of AML blood populations dependent of the AML-MSCs presence in the microenvironment.

Importantly, we reported for the first time the isolation and sequencing of the MSC fraction from autologOss. Although currently limited by low cellular yield and thus remaining a proof of concept, our analysis identified a stromal subset with high *CXCL12* expression, consistent with previously described CXCL12-abundant reticular (CAR) cells [32]. Notably, we detected a clear proinflammatory signature in AML-MSCs from autologOss, in line with scRNA-seq studies of whole patient BM [33], highlighting the ability of our model to recapitulate this feature. In contrast, our *in vitro* expanded AML-MSCs from the patient sample lost this inflammatory profile. A similar phenomenon has been described in *in vitro* cultured MDS-MSCs, which requires re-exposure to leukemic cells to reversibly regain an inflammatory state [34].

Furthermore, the combined sequencing of MSCs and hematopoietic cells enables, for the first time, interaction mapping in an *in vivo* setting. This revealed that AML-MSCs act as an interaction hub, engaging extensively with key hematopoietic populations, including the HSC/MPP compartment which likely contains leukemic stem cells. Such analyses provide a framework to identify critical communication axes that could be targeted to disrupt AML engraftment or proliferation. Expanding this approach to patients with diverse cytogenetic backgrounds will be essential to fully characterize AML to MSC interactions within autologOss and uncover novel BM niche mechanisms driving leukemia progression.

Despite clear identification of an MSC cluster, subtypes and other anticipated progenies could not be retrieved or sequenced. These include human osteoblasts, adipocytes, and potential stromal subclusters previously identified to compose hOss [35]. Their absence may be explained by the well-known challenge of retrieving MSC populations from digested bone, as well as their poor survival in microfluidic systems [36,37]. Optimization of hOss processing, together with a higher number of MSCs, would be key to identify patient-specific patterns of niche composition and their influence on disease evolution. As an alternative, spatial omics would represent a less destructive method, as well as highly multiplexed imaging methods such as CODEX (codetection by indexing), enabling simultaneous detection of dozens of protein markers at single-cell resolution while maintaining spatial relationships within intact BM niches [38,39]. Applied to autologOss, these approaches will reveal rare or spatially restricted MSC subpopulations and reconstruct interactions between stromal cells and AML blasts within defined niches. However, their use in bone-derived or mineralized tissues remains technically demanding and requires further optimization for reliable tissue processing and signal resolution.

Last, novel 3D *in vitro* culture systems have emerged as an alternative to recapitulate the human BM niche, particularly those based on iPS cells [37,40,41]. While promising, these systems have not been validated for AML culture. Besides, concerns remain on whether they can faithfully reproduce key BM niche features and functions [19], as they rely on developmental differentiation program leading to fetal hematopoiesis [40,41]. However, they present the advantage of being fully human and highly controllable systems, whereas hOss are inherently chimeric. In contrast, hOss lead to a more diverse and physiological recapitulation of the mesenchymal elements of the BM, and the functional maintenance of leukemia.

To conclude, our study proposes OssiGel as a new technology enabling the unprecedent formation of autologOss. The generation of standardized-yet patient-specific-ossicles offers a promising platform for functional drug testing. This model may allow more predictive assessment of therapeutic responses and resistance mechanisms, supporting individualized treatment strategies. Ultimately, it represents a step toward integrating personalized disease modeling into clinical decision-making.

## Acknowledgements

We gratefully acknowledge the LBIC (Lund Bioimaging Centre), the CTG (Center for Translational Genomics), the Lund Stem Cell Center FACS facility, the Lund Stem Cell Center imaging facility, the CAGT (Cell and Gene Technologies Core), and the CCM (Centre for Comparative Medicine) from Lund University. This work was supported by Cancerfonden (to P.B.), European Research Council (Starting grant #948588 and Proof-Of-Concept to P.B.), the Knut and Wallenberg foundation (Wallenberg Launch Pad, to P.B), Vinnova (Innovation Stage 1, to P.B.), Region Skåne and Faculty of Medicine at Lund University.

## Authors contribution

C.S. Experimental design, performing experiments, data visualization, writing – original draft; D.Z. Experimental design, performing experiments, writing – review and editing; A.G.G. Experimental design, performing experiments, data visualization, writing – Original draft; D.H.G. Single-cell sequencing data analyses and visualization, writing – original draft; Y.L. Single-cell sequencing data analyses and visualization; A.G. Conceptualization, performing experiments; M.G. Performing experiments; A.B. Performing experiments, writing – review and editing; S.G.A. Performing experiments, writing – review and editing; J.C. Resources (healthy donor BM samples); V.L. Resources (AML patient BM samples); P.B. Conceptualization, experimental design, supervision, funding acquisition, writing – review and editing.

## Materials and Methods

### MSOD-B

MSOD-B cells (P21-P22) were expanded in complete medium consisting of alpha Minimum Essential Medium (aMEM) supplemented with 10% fetal bovine serum, 1% HEPES (1M), 1% sodium pyruvate (100mM), 1% of penicillin/Streptomycin/Glutamine (100X) solution and 5 ng/ml of FGFb (all purchased from Gibco, USA) under standard culture conditions until confluence (90%). Medium was replaced twice a week. Cells were seeded at a density of 3,400 cells/cm^2^.

### MSOD-B differentiation on collagen scaffolds

MSOD-B cells (P23) were harvested using trypsin (Gibco, USA) and seeded on cylindrical type I collagen sponges (Avitene^TM^ Ultrafoam^TM^ Collagen Sponge, Davol, USA) of 6 mm diameter and 3 mm thickness. Collagen scaffolds are shaped with a 6 mm biopsy punch (Kai Medical, USA) and placed in a well of a 12-well plate coated with 1% agarose to prevent cell adhesion. Per scaffold, 35µL of cell suspension containing 2M of cells were seeded at the surface and incubated 1h under standard culture conditions (MSOD-B expansion medium without FGFb). Then 2mL of chondrogenic medium is added to each well. Medium is composed of DMEM, 1% Insulin/Transferrin/Selenium, 1% sodium pyruvate (100mM), 1% of Penicillin/Streptomycin/Glutamine (100X) solution (all from Gibco Invitrogen, USA), 0.47 mg/mL linoleic acid (Sigma-Aldrich, USA), 25 mg/mL bovine serum albumin (Sigma Aldrich), 0.1 mM ascorbic acid-2-phosphate (Sigma-Aldrich), 10 ng/ml TGF-b3 (R&D Systems, USA) and 0.1 μM dexamethasone (Sigma-Aldrich).

### Lyophilization

After three weeks of differentiation, MSOD-B constructs are rinsed twice with PBS pH 7.2 (Gibco, USA). For lyophilization, constructs are then snap froze inside 15 ml tubes in liquid nitrogen for 5 minutes and then lyophilized at-80°C and 0.024 mbar overnight (Labconco, USA). Once freeze-dried, the lyophilized cell constructs are stored at 4°C for long-term storage.

### Powder generation

Lyophilized cartilages were ground with the CryoMill (Retsch, Germany) in a 10 ml stainless-steel grinding jar containing two 10 mm diameter grinding spheres. To ensure that the samples were pre-embrittled before grinding, a pre-cooling time of 12 minutes was applied. Grinding was then performed four times for 2 minutes at 25 Hz, with intermediate cooling periods of 30 seconds at 5 Hz. The resulting powder was stored at 4°C for long-term storage.

### SEM on powder

The powder samples were sputter-coated with a 10 nm layer of platinum/palladium (80:20 ratio), mounted on stubs, and examined using a JEOL JSM-7800F field emission scanning electron microscope (FEG-SEM) at the Lund University Bioimaging Centre (LBIC).

### GAG quantification

After lyophilization, engineered human cartilages were digested overnight at 56°C in 0.5 mL of a proteinase K solution (1 mg/mL proteinase K; 10 μg/mL pepstatin A; 1 mM EDTA; 100 mM iodoacetamide; 50 mM Tris, all from Sigma-Aldrich) at pH 7.6. Samples were dissolved by vortexing and diluted 1:5 with distilled water. GAGs were quantified using the Blyscan Sulfated Glycosaminoglycan Assay (Biocolor Ltd, United Kingdom), following the manufacturer’s recommendations.

### AML patients BM samples

Fresh BM aspirates from AML patients were obtained under written informed consent in accordance with the Declaration of Helsinki under the following ethical approvals: 2021-04046 (healthy donor BM samples) and 2021-00947 (AML patient BM samples). Human CD3^+^ cells were depleted using RosetteSep (#15661, StemCell Technologies, Canada) or a MACS depletion column (#130-097-043, Miltenyi Biotec) following the manufacturer’s recommendations. BM aspirates were filtered through a 70 mm strainer. The samples were diluted 1:2 with MACS buffer. Then, density gradient centrifugation was performed using LMS 1077 Lymphocyte (PAA Labs, Austria) to isolate total mononucleated cells, at 300xg for 30 min with breaks off. Interphase was collected and diluted with cold MACS buffer. Next, red blood cells were lysed twice (#420302, Biolegend) and cells were counted using Trypan Blue. CD45 positive and negative fractions were enriched and collected separately using MACS magnetic separation (#130-045-801, Miltenyi Biotec), following the manufacturer’s recommendations.

### MSC isolation from human BM

Total mononucleated cells or CD45 negative fraction of human BM samples from AML patients or healthy donors were seeded in flasks and MSC were enriched by plastic adherence using MSC expansion medium as previously described. The medium was changed twice a week for 1-3 weeks. Once ‘colonies’ of cells were emerging or around 80% confluency, MSCs were then trypsinized and transferred into new flasks. They were cultured until P2-P4 before mixing them with OssiGel.

### Mice

NSG or FoxN1 KO BALB/C (Nude mice) (2 to 4 months old) were obtained from Charles River Laboratories. All mouse experiments and animal care were performed in compliance with the Lund University Animal Ethical Committee (ethical permits number 15485-18, 11518-20, and 9915-25) and in accordance with the 3R principles. Mice were housed in a temperature-controlled ventilated cabinet under a 12 h light-dark cycle with free access to water and a standard irradiated rodent chow diet. All mice used were maintained under specific pathogen-free conditions.

### Hydrogel formation

Four hydrogel formulations were prepared using fibrinogen sourced from either Sigma-Aldrich (F3879, Sigma-Aldrich, USA) or Baxter (Tisseel Fibrin Sealant, Baxter, USA), yielding final fibrin concentrations of 6 mg/mL and 12 mg/mL. Briefly, 2 mg of lyophilized MSOD-B powder was suspended in the respective fibrinogen solution diluted in complete medium. Once the powder was fully hydrated, thrombin (#341578, Sigma-Aldrich) was added to a final concentration of 1.5 U/mL and mixed thoroughly. The hydrogels were allowed to reticulate for 30 minutes at 37°C prior to *in vitro* use. For *in vivo* injectability assessments, formulation 3 (Fibrin 6mg/ml, Baxter) was exclusively utilized. The hydrated powder and fibrinogen mixture was maintained at 4°C to prevent premature gelation; thrombin was added immediately prior to ectopic injection, allowing the solution to reticulate *in situ* within the animal.

### Atomic Force Microscopy (AFM)

The elastic modulus of the hydrogels was determined AFM force spectroscopy using a NanoWizard 4 system (JPK Instruments, Bruker) mounted on a Nikon Ti-2 inverted microscope with a 20× ELWD S Plan Fluor objective (NA = 0.6). Measurements were performed with rectangular cantilevers equipped with a colloidal probe of approximately 2.25 μm radius (CP-qp-CONT-Au-A, NanoAndMore), having a nominal spring constant of 0.1 N m⁻¹ and resonance frequency of 30 kHz. Indentations were conducted in PBS without calcium or magnesium, the setpoint value for all samples was 1.5 nN, the indentation speed of 2μm/s and the sampling rate was 10kHz. The Young’s modulus was estimated by applying the Hertz contact model to the resulting force-distance curves, fitting to the first 500 nm of indentations.

### Subcutaneous implantation of constructs

Hydrogels were generated *in vitro* by mixing healthy or AML-MSCs with OssiGel as previously described with formulation 3 (Fibrin 6 mg/ml, Baxter). The number of healthy-or AML patient-derived BM-MSCs contained in each construct is specified in Table 2. Hydrogels were reticulated during 30 minutes at 37 degrees. For ‘Lyo’ experiments, AML-MSCs were resuspended in 20 ul of medium and seeded on top of lyophilized constructs. The mix was then incubated for 30 minutes at 37 degrees. All constructs were kept in PBS 1X until they were implanted on the back of NSG mice (4 hydrogels per animal) under aseptic surgery conditions. Briefly, the skin was shaved, 0.5 mm openings were created at sites of implantation and subcutaneous pockets were created to fit hydrogels. Surgery was realized under anesthesia (Isoflurane 3% for induction, 2% for maintenance) and pain was relieved using the analgesic Temgesic (Vidal, France) 0.03 mg/mL, 2 µl/g weight, subcutaneously injected. Vectabond (Vector Labs, USA) was used to close the wounds.

### Microcomputed tomography (µCT)

Explanted tissues were fixed overnight with 10% formaldehyde before being subjected to *ex vivo* µCT with the U-CT system (MILabs, The Netherlands) using a tungsten x-ray source at 50 kV and 0.21 mA. For each sample, a circular scan (360°) was recorded with an incremental step size of 0.250°. Volumes were reconstituted at 10 µm isotropic voxel size using MILabs software analysis. For BV analysis, the highly mineralized tissue volume was quantified using Seg3D [v2.2.1, NIH, NCRR, Science Computing and Imaging Institute (SCI), University of Utah, USA]. For TV analysis, each sample was meshed with Blender (v2.82a, The Netherlands) and analyzed with an in-house developed script. For cortical and trabecular thickness, the measurements were performed using the SCANCO Medical software suite.

### In vivo µCT scanning

Mice were subjected to *in vivo* X-ray analysis at 4-6 weeks after surgery. Briefly, animals were anesthetized using isoflurane (3%) and placed in a right lateral decubitus position. Lower body was then scanned with a U-CT system (MILABS, Netherland) using a tungsten x-ray source at 50 kV and 0.21mA. A circular scan (360°) was recorded with an incremental step size of 0.250°. Volumes were reconstituted at 30 µm isotropic voxel size using MILABS software analysis. For BV analysis, the highly mineralized tissue volume was quantified using Seg3D (v2.2.1, NIH, NCRR, Science Computing and Imaging Institute (SCI), University of Utah, USA).

### AML-MSC transduction with TdTomato

Primary human MSCs from AML patients and healthy donors were transduced with a lentiviral vector encoding Td-Tomato to achieve stable fluorescent labeling and tracing over time. Briefly, P2 or P3 MSCs were seeded in duplicates or triplicates in a 6-well plate at 50% confluence in 1 ml of complete MSC expansion medium. Following a short incubation at 37C, the target wells were exposed to lentiviral particles with TU 10^8^/ml at MOI 10. After 12h of incubation, media were replaced from the Td-Tomato-containing wells and the control wells (only medium) with complete MSC expansion medium. The viability and expansion of Td-Tomato positive cells was monitored over time, and the transduction efficiency was assessed by fluorescence microscopy and flow cytometry. At 80% confluence Td-Tomato positive cells were FACS sorted and seeded for further expansion. In cases where the Td-Tomato signal in the transduced cells exceeded 85%-90%, cells were not sorted to avoid unnecessary cell manipulation.

### In vivo transplantation of AML cells

Between 4 and 6 weeks after OssiGel implantation, and upon successful hOss formation, all mice were sub-lethally irradiated (24h before transplantation, 200 cGy). Then, half of the mice were transplanted with AML cells either by intra-ossicle or intravenous injections. The other animals were controls, injected with PBS 1X through the same procedure. <u>Intra-ossicle injections</u>: Mice were anesthetized (Isoflurane 3% for induction, 2% for maintenance) and skin was shaved. Each hOss was maintained using forceps and a volume of 15 to 20 µl of AML cells diluted in PBS 1X (Gibco) was injected with a 500 µl syringe and a 29G needle. The number of AML cells transplanted per hOss is detailed in Table 2, and 4 hOss were injected per animal. <u>Intravenous injections</u>: Mice were held using a plastic restrainer. AML cells were diluted in PBS 1X (Gibco) and transplanted by injecting a volume of 150 µl into the tail vain. Number of AML cells transplanted per mouse is detailed in Table 2.

### Flow cytometry

Every 4 weeks, around 50 µl of blood were collected from the tail vein of the mice to follow engraftment of human AML *in vivo*. Red blood cell lysis was performed twice with ACK Lysis buffer (Thermo Fisher, USA) and then cells were washed with PBS 1X. Cells were stained in 50 µl of MACS buffer (PBS supplemented with 2 mM EDTA and 0.5% BSA) using the following antibody combination and dilutions: anti-mCD45-AF700 at 1:50 (clone HI30, #368514, Biolegend), anti-hCD45-Pacific Blue at 1:50 (clone 30-F11, #103126, Biolegend), anti-CD33-PE at 1:100 (clone WM53, #555450, BD Biosciences), anti-CD19-APC at 1:100 (clone HIB19, #302212, Biolegend), and anti-CD3-PE-Cy7 at 1:200 (clone UCHT1, #563423, BD Biosciences). Cells were incubated 20 min in the fridge, then excess antibodies were diluted in MACS buffer and removed after centrifugation (300 g, 5 min). Cells were resuspended in 300 µl of MACS buffer and 7AAD (Fisher Scientific) was used at 1:100 as a viability marker. Samples were analyzed on a flow cytometer (BD LSRFortessa or BD LSRFortessa X-20). Data was analyzed with FlowJo (version 10.10.0).

### Fluorescence-activated cell sorting

At the endpoint of experiments, mice were euthanized and the femurs, tibiae and hOss were collected. Femurs and tibiae were crushed using a porcelain mortar and pestle with PBS 1X or flushed with PBS 1X. Mortar was rinsed once with PBS 1X after crushing. Then, collected cells were filtered using a 40 µm strainer. Macroscopic pictures of the hOss were taken before and after explantation with an iPhone 15 Pro. One hOss per mouse was kept for histology and immunofluorescence staining. The remaining hOss were crushed in a 1.5 mL tube with PBS 1X and a micropestle (#10683001, Fisher Scientific). Supernatant was kept separately, and crushed hOss were cut into small pieces in digestion buffer (M199 medium (Gibco) supplemented with 1.1 mg/mL Collagenase Type II (Gibco), 0.1% BSA (Sigma-Aldrich), 100U/mL DNAse I (Invitrogen, USA), 1 mM CaCl2 (Sigma-Aldrich), 1% Poloxamer 188 (Sigma-Aldrich), 20 mM HEPES). Samples were incubated for 40-60 min at 37 degrees on an orbital shaker at 200 rpm and then filtered through a 40 µm strainer. Cold M199 medium was added to stop the digestion and the supernatant previously obtained was added. Cells were centrifuged at 300 g for 5 minutes. All organs underwent red blood cell lysis using ACK lysis buffer as previously described and then cells were counted with Trypan blue (Gibco) on an automated cell counter (TC20, Bio-Rad, USA). Cells were stained in 100 µl of MACS buffer using the following antibody references and dilutions: anti-hCD45-FITC at 1:50 (clone HI30, #304006, Biolegend), anti-mCD45-PE at 1:50 (clone 30-F11, #103126, Biolegend), anti-CD298-PE-Cy7 at 1:100 (clone S20013B, #377812, Biolegend), anti-CD73-APC at 1:200 (clone AD2, #APC-65162, Proteintech, USA), anti-CD105-APC at 1:50 (clone 266, #562408, BD Biosciences), anti-PDPN-APC at 1:50 (clone NC-08, #130-126-165, Miltenyi Biotec, Germany), anti-CD146-APC at 1:50 (clone SHM-57, #130-111-323, Miltenyi Biotec). Cells were resuspended in 300 µl of MACS buffer, and DAPI (Sigma-Aldrich) at 0.1 mg/mL was used as a viability marker. Samples were analyzed and cells were sorted on a cell sorter (BD FACSAria III). The different gating strategies are shown in supplementary figure 4E. For single-cell sequencing, cells were sorted into a 1.5 mL tube containing 200 ul of cold MACS buffer.

### Paraffin-embedding of tissues

Tissues were fixed in 4% formaldehyde (Thermo Scientific) at 4°C, for 12 h, washed three times in PBS 1X, and decalcified with 10% EDTA solution (Sigma-Aldrich), pH 8 up to 2 weeks with gentle rocking, at 4°C. Tissues were sequentially dehydrated in 35%, 70%, 95%, 99.5% graded ethanol solution (Solveco, Sweden) twice for 20 min each. Then, they were washed 20 min in 99.5% ethanol/xylene solution (1:1, Fisher Scientific) followed by two incubations of 20 min in xylene (Fisher Scientific). After immersing tissues in paraffin at 56°C overnight, they were embedded and cut with a microtome in 10 μm sections. The sections were then dried overnight at 37°C. To remove the paraffin, sections were washed with xylene for 7 min twice, then once in 99.5% ethanol/xylene solution (1:1) for 3 min. Afterwards, sections were rehydrated in 99.5%, 95%, 70%, 35% ethanol, twice for 7 min each for subsequent histological stainings.

### Safranin-O/Fast Green staining

Sections were stained with Mayer’s hematoxylin solution (Sigma-Aldrich) for 10 min. This was followed by a washing step to remove extra color from sections during 10 min under running distilled water. They were then stained with 0.01% fast green solution (Fisher Scientific) for 5-10 min, and extra color was removed by rinsing the sections quickly with 1% acetic acid solution (0.5 ml acetic acid in 49.5 ml distilled water) for no more than 15 seconds. After staining the slides with 0.1% Safranin O (Fisher Scientific) solution for 5 min, dehydration and clearing were done by immersing the slides in 95% and 99.5% graded ethanol, afterwards with 99.5% ethanol/xylene solution (1:1). The sections were finally washed in xylene twice for 2 min to remove ethanol followed by mounting with PERTEX mounting medium (HistoLab, Sweden).

### Masson’s Trichrome staining

Masson’s Trichrome staining was conducted using the Trichrome Staining Kit (Sigma-Aldrich) in accordance with the manufacturer’s protocol. Tissue sections were deparaffinized and rinsed as previously described. The sections were then placed in Bouin’s solution (Sigma-Aldrich) either overnight at room temperature or for 15 minutes at 56°C. Following this, the slides were washed under running tap water and stained with Weigert’s iron hematoxylin working solution (prepared by mixing equal volumes of solution A and B, EMD Millipore) for 5 minutes to visualize nuclei (stained black). After a wash step, the cytoplasm was stained red using Biebrich Scarlet-Acid Fuchsin for 5 minutes. The slides were then cleared by immersing them in a working phosphotungstic/phosphomolybdic acid solution (25 mL phosphotungstic acid, 25 mL phosphomolybdic acid, and 50 mL distilled water) for 5 minutes. Collagen fibers were stained blue by treating the slides with aniline blue solution for 5 minutes, followed by a clearing step in 1% acetic acid solution (prepared with glacial acetic acid, Fisher Scientific) for 2 minutes and a rinse under running deionized water. Finally, sections were dehydrated by sequential immersion in graded ethanol solutions (95% once, 100% twice) for 2 minutes each, followed by two 2-minute washes in xylene, and mounted with PERTEX mounting medium.

### Immunofluorescence staining

Samples were fixed in 4% formaldehyde (Thermo Scientific) at 4°C, for 12 h and decalcified with 10% EDTA solution (Sigma-Aldrich), pH 8 up to 2 weeks with gentle rocking, at 4°C. Embedding was performed using 4% low-melting agarose (Sigma-Aldrich) and 100 µm thick sections were cut using a 7000smz Vibratome (Campden, United Kingdom) with ceramic blades (Campden). All staining and washing steps were performed under gentle rocking in 200 µl. Sections were permeabilized using TBS 1X (Sigma-Aldrich) with 0.5% Triton X-100 (Sigma-Aldrich) for 30 minutes and then washed twice for 10 min each with TBS 0.1% Triton X-100. Sections were blocked for 2h using TBS with 10% donkey serum (Sigma-Aldrich) and 1% BSA (Miltenyi Biotec). After blocking, sections were stained overnight with primary antibodies at 4°C and washed twice with 0.1% Triton X-100 in TBS for 10 min each. Highly cross-absorbed secondary antibodies were incubated overnight at 4°C and washed twice with 0.1% Triton X-100 in TBS for 10 min each. Next, sections were incubated 10 min with DAPI at 1:100 and washed twice with TBS for 10 min each. Agarose was removed from tissue sections, and these were mounted on a slide using an antifade mounting medium (ProLong™ Diamond Antifade Mountant, Thermo Fisher). Slides were cured for 1h in a humidity chamber at room temperature and then overnight on the bench. Following primary antibodies and dilutions were used: rabbit anti-human Nuclear Antigen at 1:100 (clone 235-1R, NBP31391220UG, Novus Biologicals), rabbit anti-human CD146 at 1:100 (clone 44, #MA5-29415, Thermo Fisher), goat anti-murine CD31 at 1:100 (polyclonal, #AF3628, R&D systems, USA), goat anti-TdTomato at 1:50 (polyclonal, AB8181-200, SicGen Antibodies), rat anti-human CD45 at 1:100 (clone YTH 24.5, #MCA345GT, Bio-Rad), mouse anti-human CD33 at 1:100 (clone WM53, NBP2-32931, Novus Biologicals). The following secondary antibodies were employed: AlexaFluor488 donkey anti-rabbit (#711-545-152, Jackson ImmunoResearch, USA), Cy3 donkey anti-rat (#712-165-150, Jackson ImmunoResearch), AlexaFluor680 donkey anti-goat (#A32860, Thermo Fisher), and AlexaFluor488 donkey anti-mouse (#715-545-150, Jackson ImmunoResearch). All secondary antibodies were used at a 1:400 dilution. Samples were imaged with a Leica Stellaris confocal microscope (Leica, Germany) and images were processed using the LasX software (version 3.10.0). The LIVE/DEAD™ Viability/Cytotoxicity Kit (#L3224, ThermoFisher) was used to assess cell viability in OssiGel following the manufacturer’s protocol.

### Library preparation, sequencing and preprocessing

scRNA-seq libraries were generated using the 10x Genomics Chromium X instrument with the Chromium Single-Cell 3′ Reagent Kit v3.1 and v4 (10x Genomics), following the manufacturer’s instructions. Libraries were pooled and sequenced on an Illumina NovaSeq 6000 or NovaSeq X Plus with paired-end reads. FASTQ files were processed using Cell Ranger (v9.0.1) and aligned to the GRCh38-2020-A human reference genome.

### Single-cell data analysis

Downstream analysis of filtered feature-barcode matrices was performed in R using Seurat (v4.4.0). Quality control was applied independently to each sample prior to dataset integration. Cell type annotation was performed by reference atlas mapping using Harmony [42] and Symphony [43] to project our dataset onto the reference embedding space, followed by label transfer to assign cell type identities. Cell-cell communication analysis was performed using CellPhoneDB (v5.0) in DEG mode (method 3), based on differentially expressed genes precomputed per organ and cell type.

### Statistical analyses

Normality of data distribution was assessed with the Shapiro-Wilk test. For normally distributed data, an unpaired t-test was performed, otherwise the non-parametric Mann-Whitney test was used. Two-way ANOVA was used to analyze the difference between the means of more than two groups. All graphs and statistical analyses were performed with GraphPad Prism (v10.5.0). The statistical tests and *p* values are indicated in related figure legends, with *p* values of <0.05 considered significant.

---

**Supp. Figure 1.**
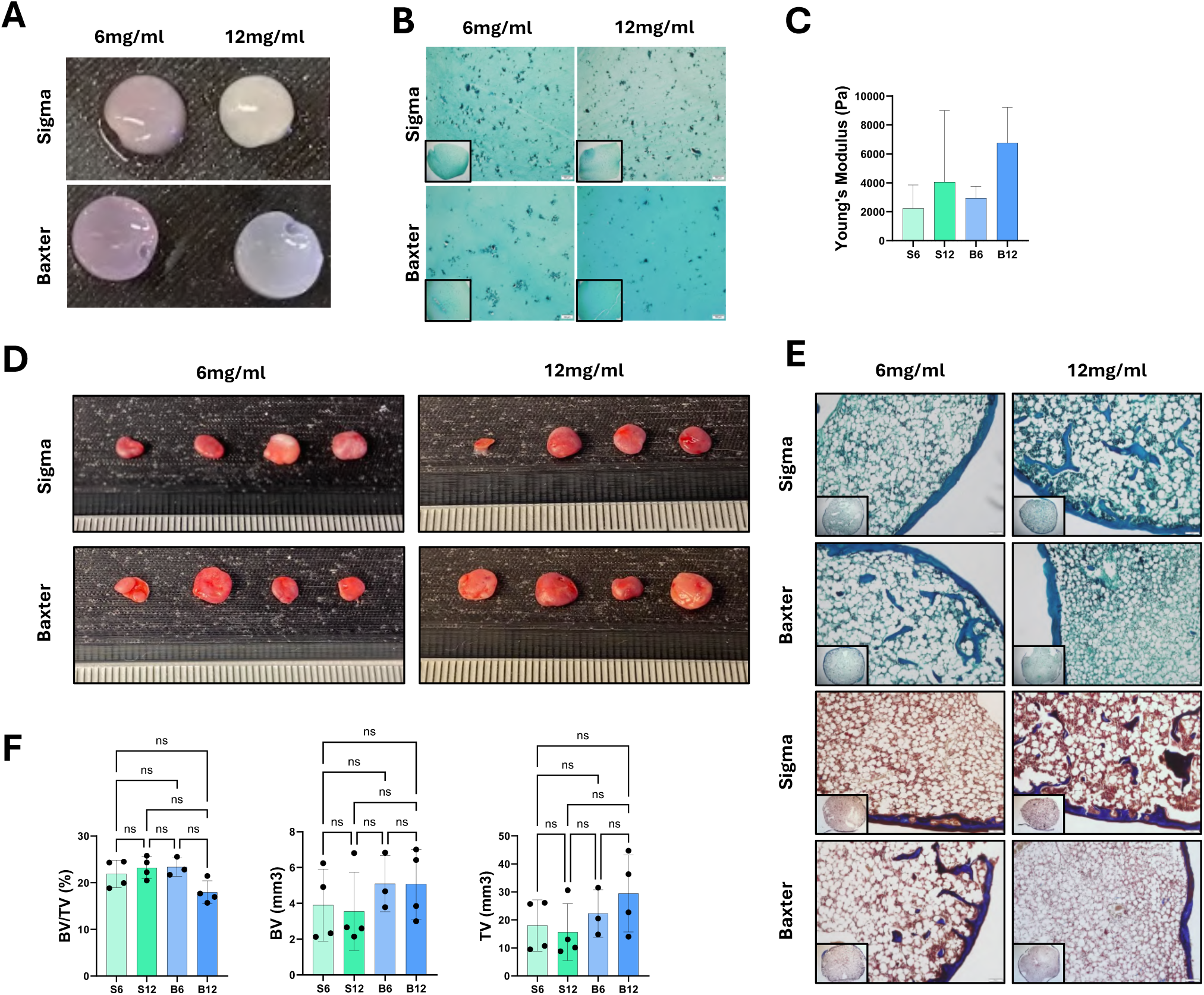
A. Macroscopic images of four different OssiGel formulations (*n*=4). **B.** Safranin-O/Fast Green staining of the different gel formulations (*n*=4). **C.** Young Modulus of each formulation assessed by atomic force microscopy (*n*=4). **D.** Macroscopic images of humanized ossicles (hOss) from different OssiGel formulations following ectopic implantation in immunodeficient mice for 6 weeks (*n*=4). **E.** Safranin-O/Fast Green and Masson’s Trichrome staining of explanted tissues after 6 weeks of ectopic implantation to assess cartilage remodeling and bone formation (*n*=4). **F.** BV/TV (%), BV (mm^3^) and TV (mm^3^) of explanted tissues 6 weeks after ectopic implantation in immunodeficient mice, assessed by µCT analysis (*n*=4). S6: Sigma 6 mg/mL, S12: Sigma 12 mg/mL, B6: Baxter 6 mg/mL, B12: Baxter 12 mg/mL.

**Supp. Figure 2.**
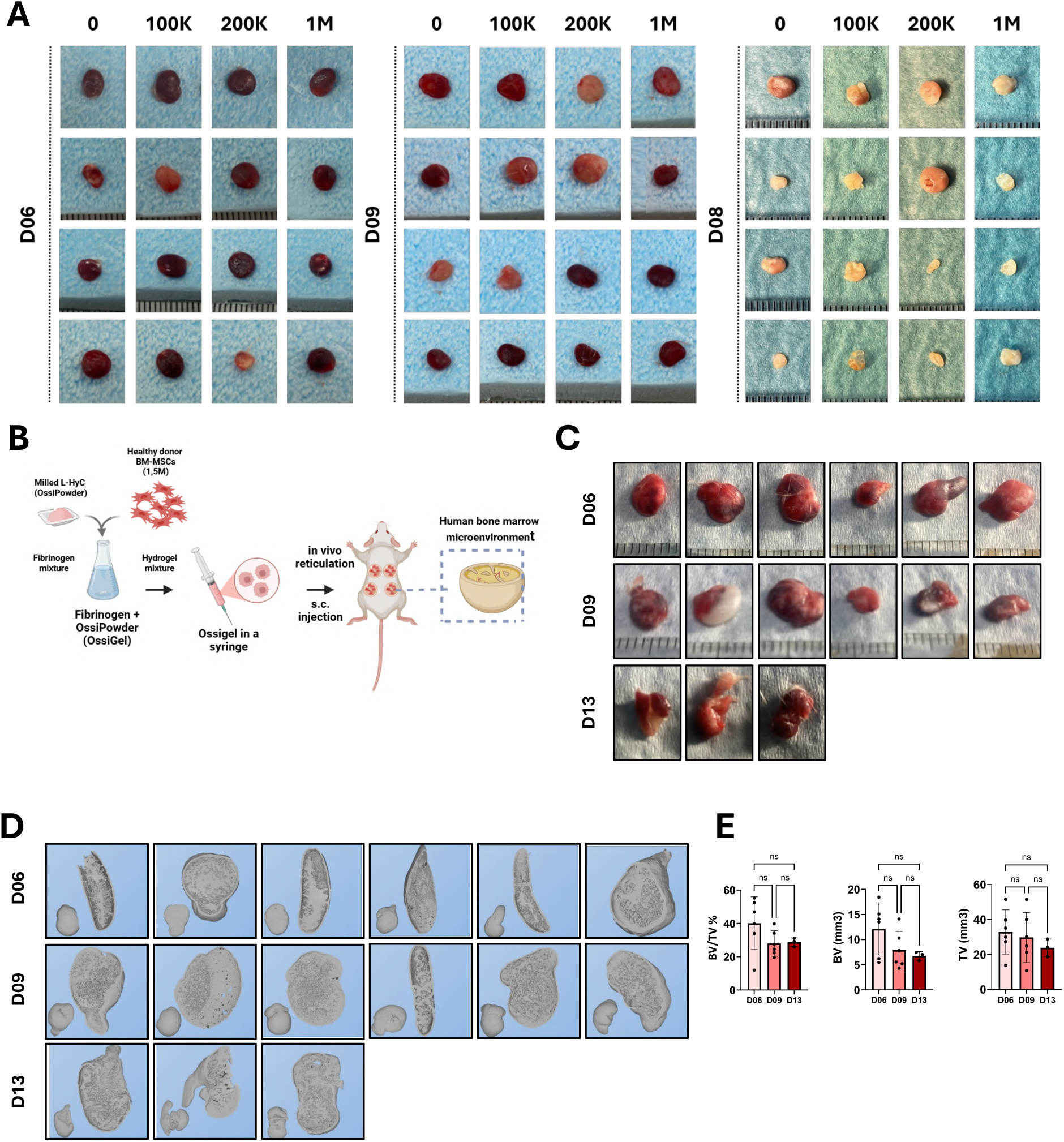
A. Macroscopic appearance of OssiGel constructs generated with increasing cell doses (0, 1×10⁵, 2×10⁵ and 1×10⁶ human donor BM-MSCs per construct) from three independent healthy donors (D06, D09 and D08) after 6 weeks of ectopic implantation (*n*=4 for each donor). **B.** Schematic overview of the experimental workflow illustrating preparation of OssiGel from milled lyophilised constructs, mixed with fibrinogen and thrombin, loaded into a syringe, and subcutaneously delivered in immunodeficient mice to generate a human bone marrow-like microenvironment. **C.** Representative macroscopic images of explanted OssiGel-based hOss obtained by injectable delivery for the three donors (D06, D09 and D13) after 6 weeks *in vivo* (*n*≥3 for each donor). **D.** Representative three-dimensional µCT reconstructions of the resulting hOss from each donor, illustrating donor-specific variability in overall shape and internal architecture (*n*≥3, 3 independent donors). **E.** Quantification of BV/TV (%), BV (mm^3^) and TV (mm^3^) of hOss derived from donors D06, D09 and D13, showing inter-donor differences in hOss size and mineralized tissue content (mean ± SD; ns, not significant, *n*≥3, 3 independent donors).

**Supp. Figure 3.**
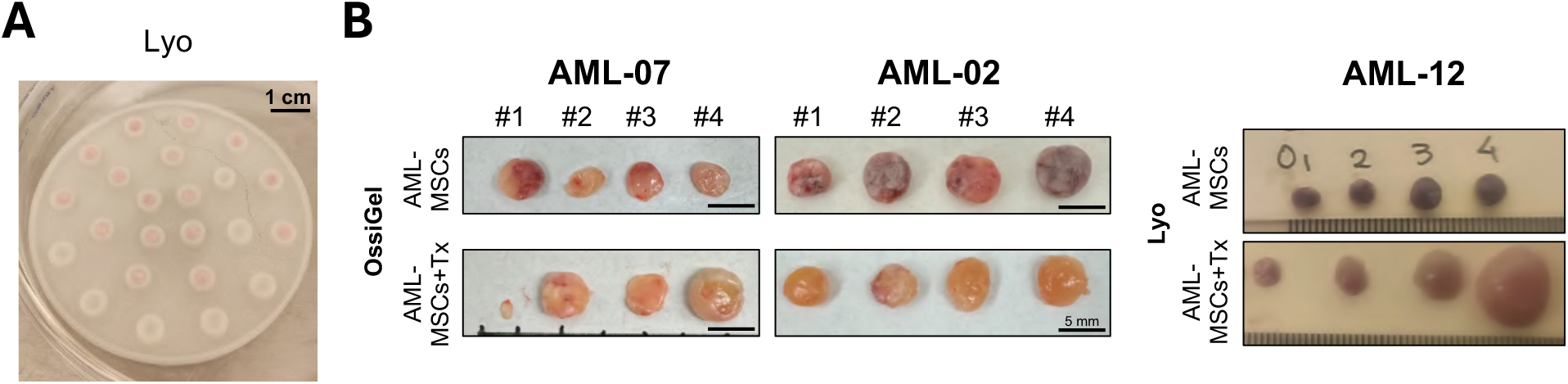
A. Macroscopic pictures of Lyo incubated with AML-MSCs before *in vivo* implantation. **B.** Macroscopic images of hOss with and without AML cell transplantation from OssiGel (AML-07, 27 weeks and AML-02, 29 weeks post-transplantation) or Lyo (AML-12, 18 weeks post transplantation).

**Supp. Figure 4.**
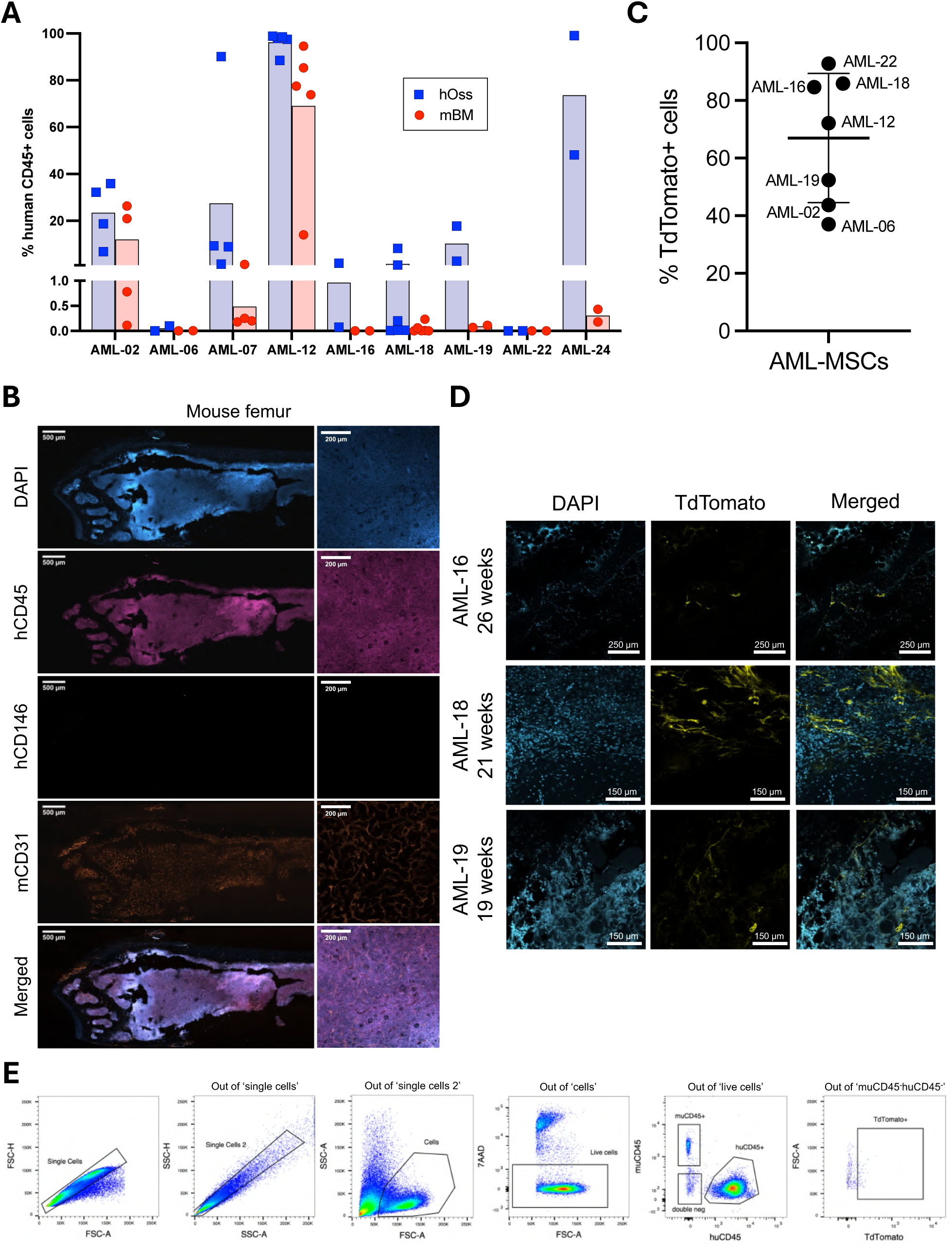
A. Percentage of hCD45^+^ cells in autologOss versus corresponding mouse femurs at endpoint of experiments. Each dot represents one animal; *n*=2 to 5 mice per patient sample. **B.** Representative confocal images of hCD45^+^ cells and CD146^+^ MSCs within mouse femur 12 weeks after transplantation. Nuclei are stained with DAPI. Maximum intensity projection of a Z-stack at magnification x10 (whole sections) or x20 (details). **C.** Lentiviral transduction efficiency of *in vitro* AML-MSCs with a lentiviral vector expressing TdTomato (MOI: 10). **D.** Long-term persistence (18-26 weeks post transplantation) of TdTomato^+^ MSCs within autologOss. Single stack images at magnification x10. **E.** Flow cytometry gating strategy applied for the Fluorescence-Activated Cell Sorting (FACS) of hematopoietic cells (huCD45^+^) and stromal cells (TdTomato^+^).

**Supp. Figure 5.**
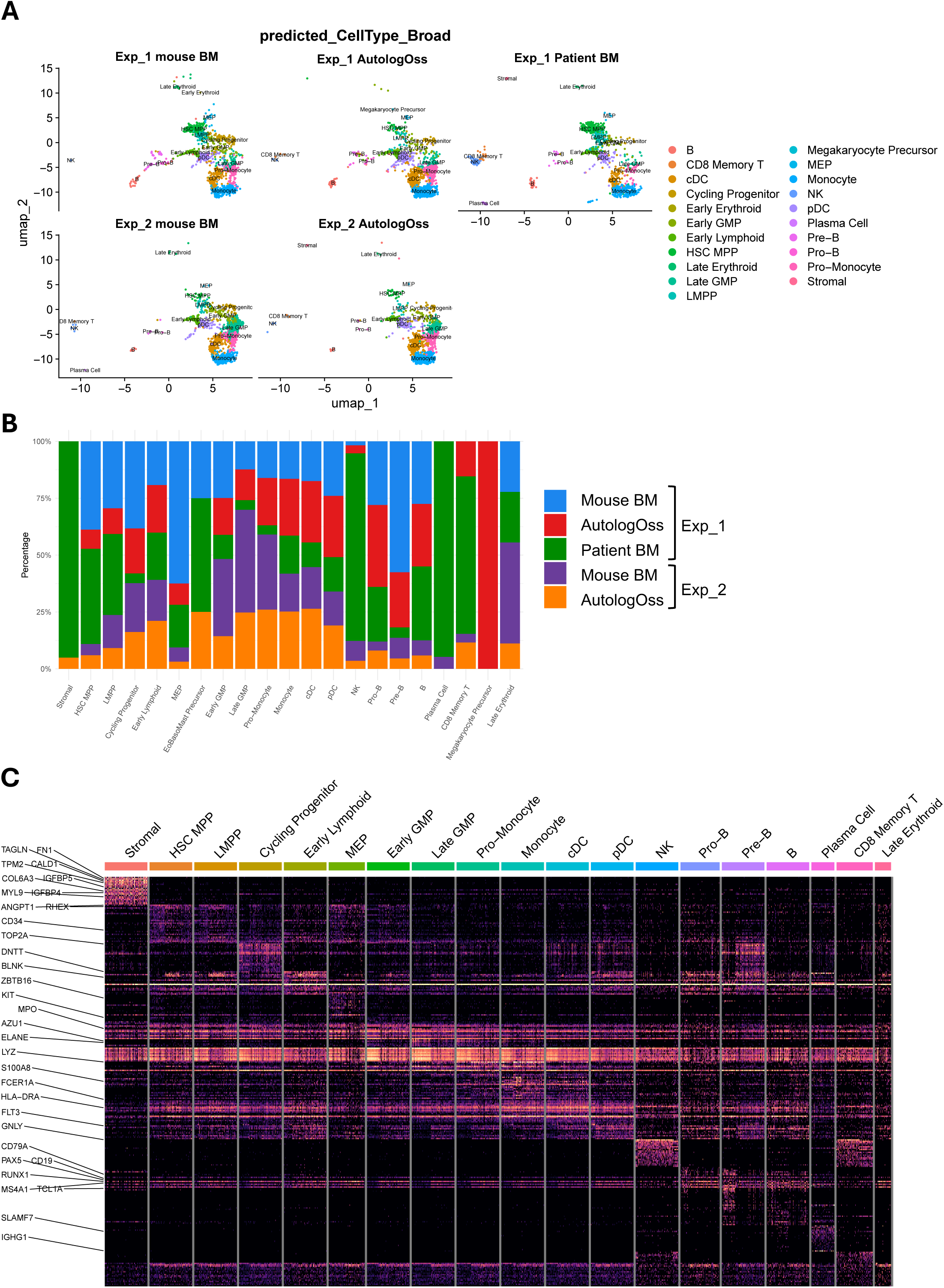
A. UMAP embedding of mapped cell types from patient BM, murine femurs and autologOss, colored by cell type. **B.** Stacked bar plot depicting the absolute or proportional distribution of individual samples within each annotated cell population. **C.** Heatmap of scaled expression values for the top differentially expressed genes per cell population, identified by test used, e.g., Wilcoxon rank-sum test. Rows represent genes; columns represent individual cells grouped and ordered by annotated cell type. Color scale indicates z-scored normalized expression.

**Supp. Figure 6.**
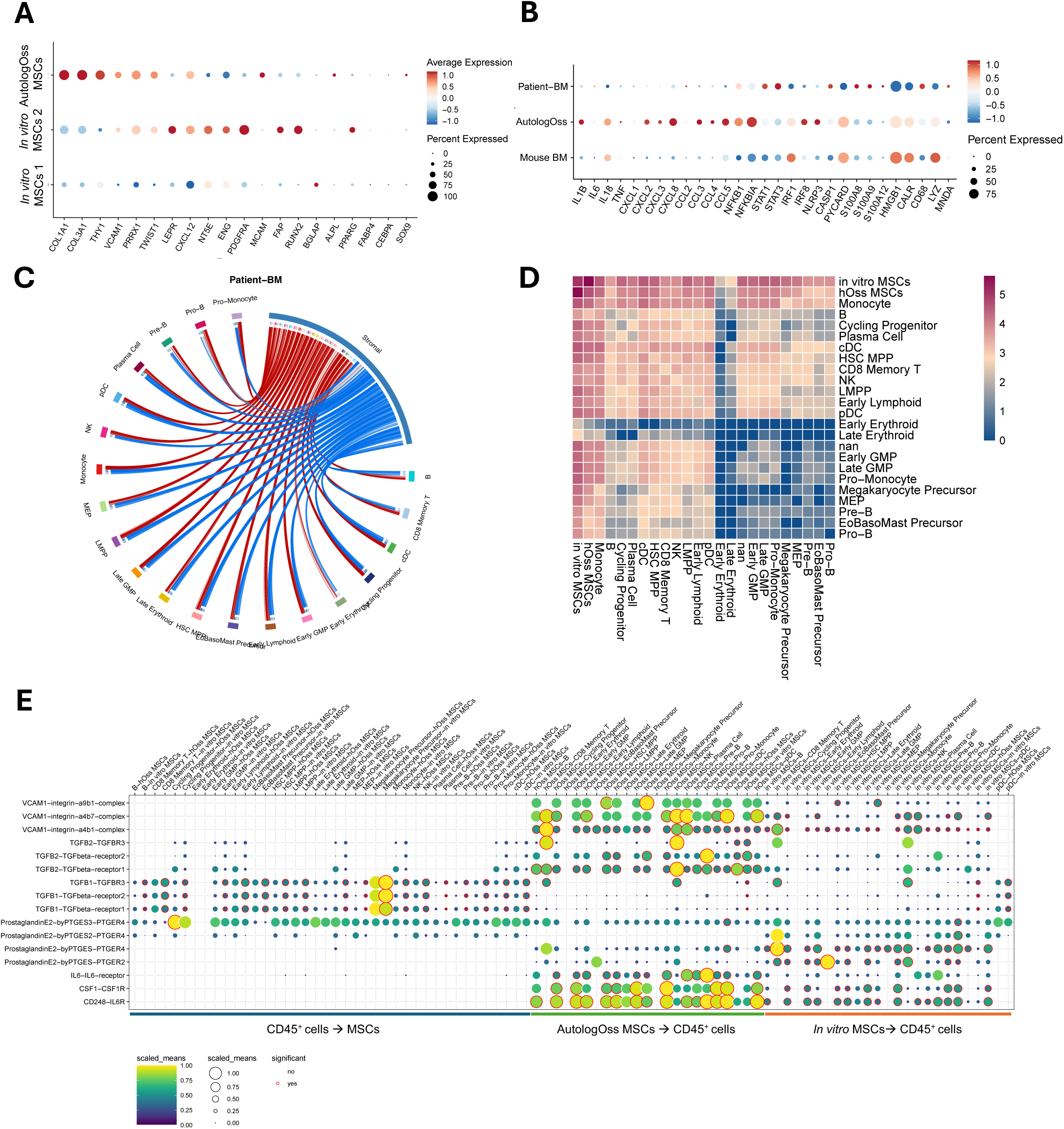
A. Dot plot showing scaled expression of inflammatory mediators in blood populations across murine, human and humanized niches. **B.** Dot plot depicting the expression of canonical MSC markers across subpopulations. **C.** Circos plot of predicted ligand-receptor interactions from patient MSCs. **D.** Heatmap displaying the number of significant ligand-receptor interactions between cell type subpopulations and all other annotated cell types, with color intensity reflecting interaction count. **E.** Dot plot providing a comprehensive overview of enriched communication pathways across all annotated cell populations and conditions, extending the pathway analysis presented in Figure 6D.

